# Nanopore translocation reveals electrophoretic force on non-canonical RNA:DNA double helix

**DOI:** 10.1101/2023.09.12.557357

**Authors:** Filip Bošković, Christopher Maffeo, Gerardo Patiño-Guillén, Ran Tivony, Aleksei Aksimentiev, Ulrich F. Keyser

**Affiliations:** Cavendish Laboratory, University of Cambridge, Cambridge, CB3 0HE, UK; Department of Physics, University of Illinois at Urbana–Champaign, Urbana, 61801, IL, USA; Beckman Institute for Advanced Science and Technology, University of Illinois at Urbana– Champaign, Urbana, 61801, IL, USA; Department of Bioengineering, University of Illinois at Urbana–Champaign, Urbana, 61801, IL, USA; Department of Chemical Engineering, Ben-Gurion University of the Negev, Beer-Sheva, 84105, Israel

**Keywords:** nanopore sensing, nanopore sequencing, RNA DNA hybrid, RNA nanotechnology, DNA nanotechnology, MD simulations

## Abstract

Electrophoretic transport plays a pivotal role in advancing sensing technologies. So far, systematic studies have focused on translocation of canonical B-form or A-form nucleic acids, while direct RNA analysis is emerging as the new frontier for nanopore sensing and sequencing. Here, we compare the less-explored dynamics of non-canonical RNA:DNA hybrids in electrophoretic transport with the well-researched transport of B-form DNA. Using DNA/RNA nanotechnology and solid-state nanopores, the translocation of RNA:DNA (RD) and DNA:DNA (DD) duplexes was examined. Notably, RD duplexes were found to translocate through nanopores faster than DD duplexes, despite containing the same number of base pairs. Our experiments reveal that RD duplexes present a non-canonical helix with distinct transport properties from B-form DD molecules. We find RD and DD molecules with the same contour length move with comparable velocity through nanopores. We examined the physical characteristics of both duplex forms using atomic force microscopy, atomistic molecular dynamics simulations, agarose gel electrophoresis, and dynamic light scattering measurements. With the help of coarse-grained and molecular dynamics simulations, we find the effective force per unit length applied by the electric field to a fragment of RD or DD duplex in nanopores with various geometries or shapes to be approximately the same within experimental errors. Our results shed light on the significance of helical form in nucleic acid translocation, with implications for RNA sensing, sequencing, and molecular understanding of electrophoretic transport.

## INTRODUCTION

Molecular transport encompasses the movement of molecules across various mediums. One specific type is electrophoretic transport, where charged molecules migrate in response to an external electric field. ^1,2^ Such electrophoretic transport is central to nucleic acid analysis, particularly in gel electrophoresis, where DNA or RNA fragments are separated according to their size by the degree of their displacement through a gel matrix under the influence of an electric field. The electrophoretic transport is paramount to nanopore-based technologies where individual nucleic acid molecules are captured and driven through a nanopore by an external electric field, enabling rapid molecular sensing. ^3–8^

Nanopores have emerged as a versatile tool to study electrophoretic transport of nucleic acids at the single-molecule level. ^3,7,9–12^ Previous studies have primarily focused on understanding nanopore translocation of DNA:DNA (DD) and RNA:RNA (RR) duplex molecules. ^13–15^ The key physical quantity governing such transport is the effective force (*F_E_*), that the external electric field, *E*, applies on the translocating molecule, conditioning both the capture rate and the translocation velocity of the molecule. ^15,16^ The nanopore translocation is opposed by the drag force of the solvent (*F_d_*) that, for translocations at low Reynolds number, is equal in magnitude and opposite in direction to the driving force. Thus, the electrophoretic transport of polyanionic polymers is primarily governed by the balance of the drag and electrophoretic forces.

Numerous factors can influence nucleic acid duplex translocation through a nanopore. One of the most studied factors that affects the nucleic acid velocity is the charge of the nucleic acid. ^14,16–18^ At physiological pH conditions, nucleic acid duplexes carry a negative charge because of their phosphate backbone, which ensures unidirectional translocation through a nanopore under external electric field. ^13,19–21^ The length of the duplex was found to affect the translocation velocity through friction with the solvent even outside of the nanopore, with longer duplexes experiencing higher drag forces and hence slower translocation rates compared to shorter duplexes. ^22^ Furthermore, the diversity of nanopore shapes and materials can contribute additional factors that affect the translocation behavior. ^9,19,22–25^ Understanding and manipulating all of these factors are essential for controlling and optimizing nucleic acid duplex translocation dynamics in nanopore-based applications.

Herein, we demonstrated how different forms of a nucleic acid duplex, namely B-form DD duplexes and non-canonical RNA:DNA (RD) duplexes. ^26–28^ influence their electrophoretic velocity in nanopore translocation experiments. Given the increasing interest in nanopore detection of RD constructs, ^29–36^ we characterize here the factors affecting the translocation of RD molecules using equivalent data for DD molecules as a reference. We find that expressing the length of the duplex as a total base pair count or as its contour length has a major effect when comparing the translocation kinetics of RD and DD molecules. We determine the molecular causes of our experimental observations through a combination of gel electrophoresis, atomic force microscopy (AFM), dynamic light scattering and molecular dynamics (MD) simulations. MD simulations and AFM imaging confirmed that RD is a A-form-like duplex. We show that the effective force acting on a unit length of A-form-like and B-form duplexes in a nanopore is nearly the same and that the difference in the translocation kinetics of the mixed duplex or B-form duplexes can be attributed to the difference of their contour lengths. We show that the persistence length and nanopore geometry have a negligible influence on the effective force. These insights are expected to facilitate accurate analysis of RNA structural isoforms and contribute to the broader understanding of electric field-driven translocation through nanoscale constrictions.

## RESULTS

### RD duplex translocates faster through a nanopore than the B-form duplex with the same number of base pairs

In our experiment, we investigated the nanopore translocation of nucleic acid duplexes using RNA or DNA as scaffolds for short complementary DNA oligos (**Figures 1a**, the design of RD and DD structures are shown in **Figure S2** and **S3**, respectively). We utilized MS2 RNA, which is 3,569 nt long, as the RNA scaffold. Initially, single-stranded M13 DNA, with a length of 7,249 nt, was double-cut using two restriction endonucleases to create a DNA scaffold constituted by a similar number of base pairs (DraIII and BaeGI; the Supporting Information **Figure S1a-c**). As an example, the double-cut M13 DNA produced a DNA fragment 3,621 nt long (**Figure 1**), which differs only in 0.2% from the length in base pairs of MS2 RNA. The DNA oligos used for assembling the 3.6 kbp RD and DD duplexes can be found in Tables S1 and S2, respectively. The reason to choose RNA and DNA scaffolds, each 3.6 kbp in length, is to allow for duplex identification when analyzed concurrently in a single nanopore measurement.

**Figure 1.**
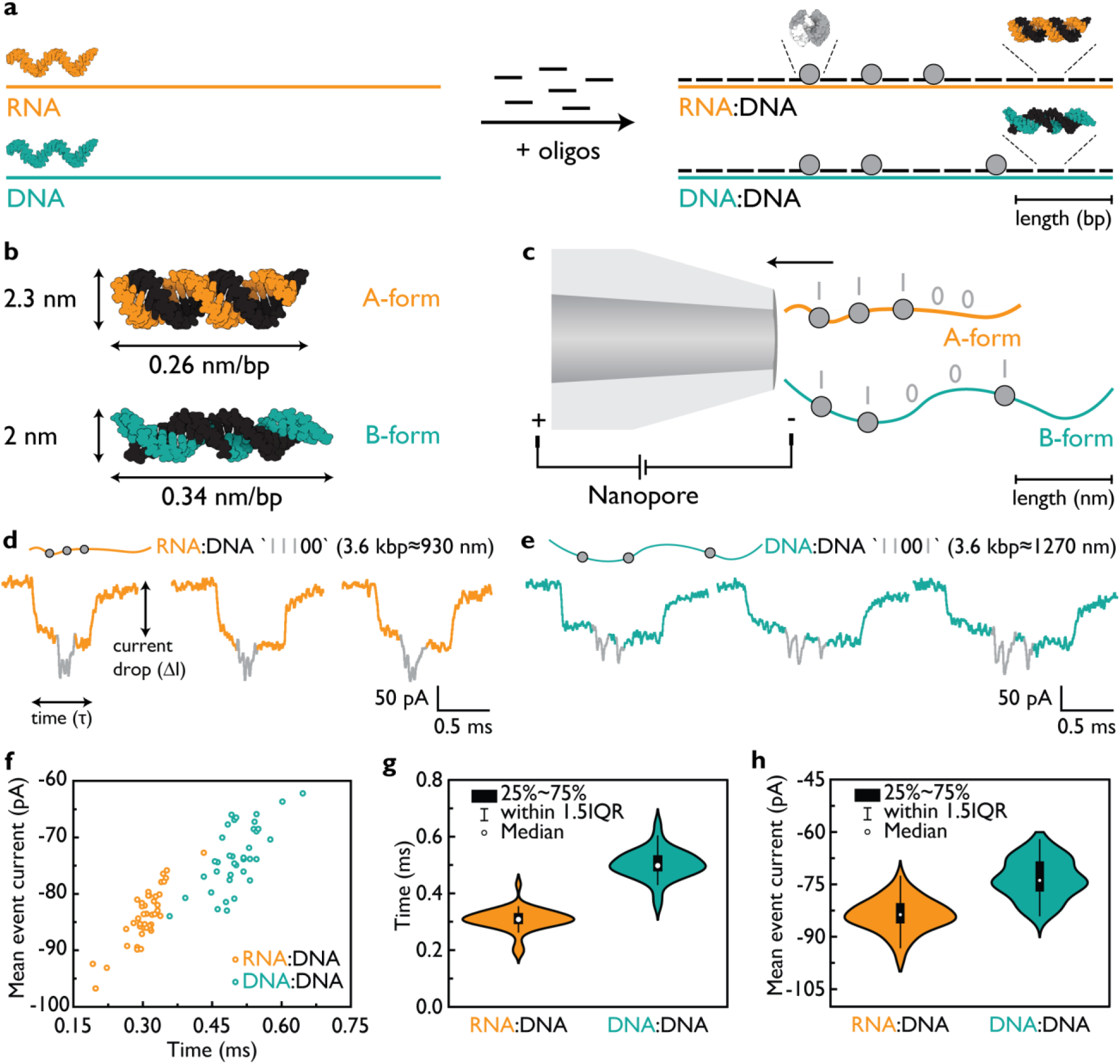
RNA:DNA (RD) and DNA:DNA (DD) duplexes with the same number of base pairs exhibit different velocities. (**a**) Single-stranded RNA (orange) and DNA (turquoise) molecules, approximately 3.6 kbp in length were hybridized with short DNA oligonucleotides (25-38 nt; black). The assembly of RD (orange:black) and DD (turquoise:black) molecules is schematically presented. Biotinylated oligonucleotides (grey circles) were strategically placed at specific positions, enabling binding with monovalent streptavidin (white region indicates biotin-binding domain) to generate a molecule-specific code for distinguishing RD and DD in nanopore measurements. RD produces the ID ‘11100’ since grey labels are equally interspaced, and DD produces ID ‘11001’ where ‘0’ indicates the absence of a label. (**b**) Proposed model structures of RD and DD duplexes, corresponding to A-form-like and B-form, respectively. The A-form-like corresponds to the shorter and wider RD. (**c**) Electrophoretically driven negatively charged RD or DD duplexes translocate through the nanopore toward the positively charged electrode. Example nanopore events of RD and DD duplexes are depicted in (**d**) and (**e**) respectively. Grey downward spikes indicate labels and the ID as ‘11100’ for RD and ‘11001’ for DD. (**f**) A scatter plot of the mean event current versus τ is shown for both RD and DD. The plot indicates two distinct populations corresponding to RD and DD codes ‘11100’ and ‘11001’, respectively. The experiments were repeated with three different nanopores for both RD and DD present together in the solution. (**g**) The translocation time (τ) of RD is 1.80 ± 0.21 times shorter than that of DD of the similar number of base pairs. (**h**) The current blockage (ΔI) of the RD duplex is 1.14 ± 0.01 times lower than that of DD. The sample size was 80 linear nanopore events for f-h. The errors represent ± standard error of the mean.

**Figure 1b** displays the physical characteristics of RD and DD duplexes. RD is expected to form an A-form-like duplex with a base pair rise of approximately 0.26 nm and a width of 2.3 nm. In contrast, DD forms a canonical B-form duplex with a base pair rise of approximately 0.34 nm and a width of 2 nm. Consequently, the contour length of RD is 24% shorter than that of DD for the same number of base pairs. Both polymers are translocated through the same nanopore (**Figure 1c**) to assess current blockage, translocation time, and determine their velocities independent of nanopore geometry and electric field distribution.

Nanopore translocation time refers to the duration required for a polymer, such as a nucleic acid duplex, to pass through the nanopore from one end to the other, driven by an electrophoretic force. ^9,19^ Consequently, by monitoring the ionic current over time, we can observe a drop in current, indicating the translocation of the duplex. In our case, the wider duplex, RD, is expected to induce a larger current drop, while the longer DD should block the ionic current for longer.

Another characteristic that we monitor during duplex translocation is the event charge deficit (ECD) which represents the area of a nanopore event. Previous studies have demonstrated that DNA can translocate in a folded conformation, ^24^ resulting in events with similar ECD compared to linear conformation events, albeit shorter and deeper. Therefore, one might hypothesize that the shorter and wider RD would exhibit a comparable ECD to the longer and thinner DD.

We conducted measurements of RD and DD samples using glass nanopores with a diameter of approximately 10 nm ^37^ (**Figure 1d-h**; nanopore details are given in Table S3). The RD identifier (ID) was designed as ‘11100’ and the DD ID was designed as ‘11001’ (**Figure 1d and 1e**, respectively; Figure S2 and S3) with nearly identical molecular weight. The different IDs allowed us to discriminate between RD and DD, enabling simultaneous nanopore measurement of both species. In the ID sequence, a ‘1’ represents a site with a 3’ biotinylated oligonucleotide that can bind with monovalent streptavidin and induce additional current blockage for molecule identification, while ‘0’ represents a site without the biotin-streptavidin conjugate. The base pair distance between neighboring ‘1’ and ‘0’ was the same (**Figure 1a**). Example nanopore events show the ‘11100’ ID readout for RD and the ‘11001’ ID readout for DD, as shown in **Figure 1d and e**, respectively (more example events are provided in **Figure S4**).

We collected data on ECD, translocation time, and current drop for RD and DD nanopore events (**Figure 1f, g, h**; additional independent measurements are provided in **Figure S5**). RD translocates close to approximately 1.80 ± 0.21 times faster than DD (for three different nanopore measurements 1.71 ± 0.12 times faster), with translocation times of *τ_RD_* ≈ 0.32 ± 0.01 ms and *τ_DD_* ≈ 0.57 ± 0.01 ms for RD and DD, respectively (**Figure 1f, g**). As expected, RD exhibits a deeper current blockage due to its wider diameter (**Figure 1h**), however this does not compensate for the shorter translocation time in the ECD ^38^ meaning that the velocity is not identical. Further confirmation is provided by the mean event current versus time for all events that clearly indicates two main non-overlapping populations (**Figure 1f**). To delve deeper into this unexpected difference in translocation times, we designed RD and DD duplexes with similar contour lengths.

### Contour length of a nucleic acid duplex governs translocation velocity rather than the number of base pairs

We aimed to investigate whether the contour length, rather than the number of base pairs, plays a significant role in the translocation velocity of a nucleic acid duplex through a nanopore. To this end, we created a 2.7 kbp DNA fragment (DD) with an absolute length of approximately 930 nm, matching the contour length of the 3.6 kbp RD duplex (930 nm). The 2.7 kbp single-stranded DNA was generated by cutting a 7,249 nt ssM13 with specific restriction enzymes (DrdI and AfeI; **Figure S1d-f**) and used as a scaffold to assemble the DD duplex with the ID ‘11001’ (the design of DD is shown in **Figure S6;** the sequences for 2.7 kbp DD are listed in Table S4). Again, we performed parallel measurements of both RD and DD constructs using multiple nanopores to observe their translocation behavior, as shown in **Figure 2** (additional independent measurements are provided in **Figure S7**).

**Figure 2.**
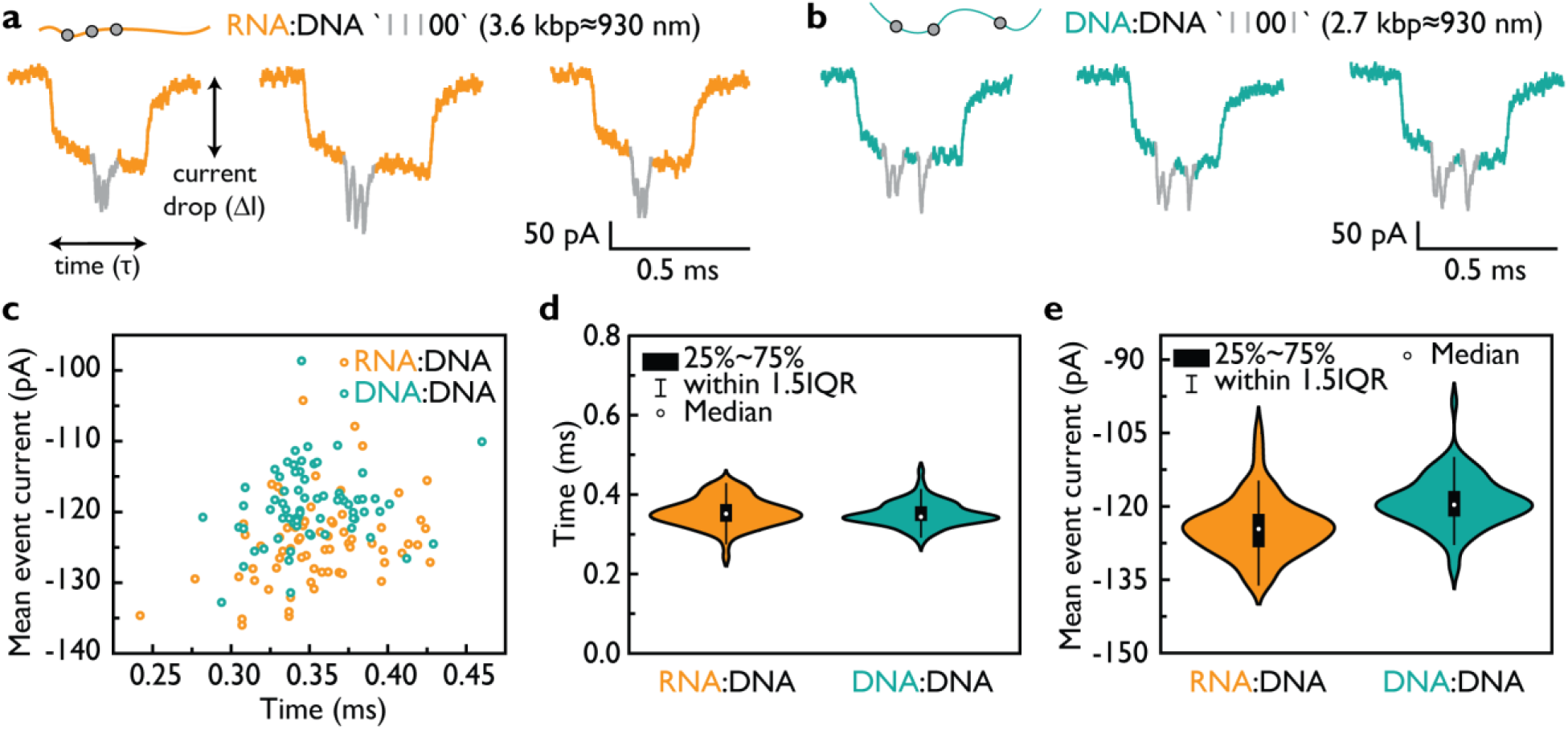
RD and DD duplexes with the same contour length exhibit comparable translocation velocity. We generated 3.6 kbp RD and 2.7 kbp DD molecules with the same absolute length of 930 nm and found that they translocate through the nanopore at similar velocities. Example events of 3.6 kbp RD and 2.7 kbp DD duplexes are presented in (**a**) and (**b**) respectively. Grey downward spikes indicate labels and the ID as ‘11100’ for RD and ‘11001’ for DD. (**c**) A scatter plot of the mean event current versus τ is shown for both RD and DD. The plot indicates overlapping populations corresponding to RD and DD codes ‘11100’ and ‘11001’, respectively. The experiments were repeated with multiple nanopores for both RD and DD present together in the solution. (**d**) The translocation time (τ) of RD 0.36 ± 0.01 ms, is similar to that of DD 0.35 ± 0.01 ms with the similar contour length. (**e**) The current blockage (ΔI) of the RD duplex is larger magnitude than that of DD. The sample size was 138 linear nanopore events for c-e. The errors represent ± standard error of the mean.

Example events of RD ‘11100’ and DD ‘11001’ are presented in **Figure 2a** and b, respectively (additional events are displayed in **Figure S8**). It is noteworthy that their translocation velocities appear similar despite RD having 25% more base pairs than DD. We plotted the translocation time of the RD and DD duplexes, revealing translocation times of 0.36 ± 0.01 ms and 0.35 ± 0.01 ms and hence similar velocities (**Figure 2c, d**). As in the initial experiments (presented in **Figure 1**), the mean current drop of RD is higher than that of DD due to its wider structure (**Figure 2e**). The scatter plot demonstrates that RD and DD events exhibit similar translocation times, while RD displays a higher current drop (**Figure 2c**). Therefore, we can conclude that the absolute length of the duplex, rather than its overall charge or molecular weight, is the primary factor determining the velocity of the duplex during translocation. The absolute values of the translocation time and of the mean event current in **Figure 2** can be different from **Figure 1** due to the use of different nanopores with slightly varying opening angle and diameter (nanopore details are given in Table S3 and S5). Hence, we repeated the measurements with at least three different nanopores to confirm that the ratio between RD and DD is consistent for the RD and DD duplexes of the same contour length (**Figure S7**). Lastly, we verified that the translocation time of RD and DD does not alter significantly during the nanopore measurement (**Figure S9**).

Lastly, we confirmed that nicks or the labels do not explain our observations. We created a nick-free DD with a sequence identical to DD ‘11001’ (for more details see **Section 3** of the Supporting Information and **Figure S10**). The translocation times for the 3.6 kbp DD with 99 nicks and without any nicks were both 0.58 ± 0.04 ms (see Figure S10c).

### Length and zeta potential measurements verify the expected values for RD and DD

In order to validate the physical characteristics for RD and DD, we conducted length and zeta potential measurements. Initially, we determined the length of the duplexes in base pairs using agarose gel electrophoresis (**Figure 3a**). Lane 1 of the gel represents a 1 kbp ladder (NEB) for reference. Lanes 2 and 3 correspond to the MS2 RNA and cut M13 DNA scaffolds, respectively. The cut M13 scaffold generates two single-stranded fragments of approximately 3,600 nt in length, with the top band representing a single-cut portion of ssM13 (7,249 nt). The assembled DD duplexes in different salt solutions are displayed in lanes 4 and 5, while assembled RD duplexes are presented in lanes 6 and 7. It is evident that both RD and DD duplexes are efficiently assembled in both 10 mM MgCl_2_ and 100 mM LiCl. In the case of DD assembled in 10 mM MgCl_2_, some structures appear to partially aggregate, whereas this aggregation is not visible in the DD duplexes assembled in LiCl solution. The presence of multiple bands in DD is expected due to the presence of three sequences: single-cut linear M13 (7,249 nt) and two fragments produced by double-cut M13 (≈3,600 nt). The majority of molecules correspond to DD (≈3,600 bp in length) with ID ‘11001’, and the bottom band corresponds to the second half of the cut M13 (≈3,600 nt) denoted as D’. The topmost band represents uncut M13, where one half is hybridized with the ‘11001’ ID oligos (DD) and the other half remains unpaired (D’). The agarose gel indicates that these two duplexes have similar molecular weight *i.e.* the similar number of base pairs.

**Figure 3.**
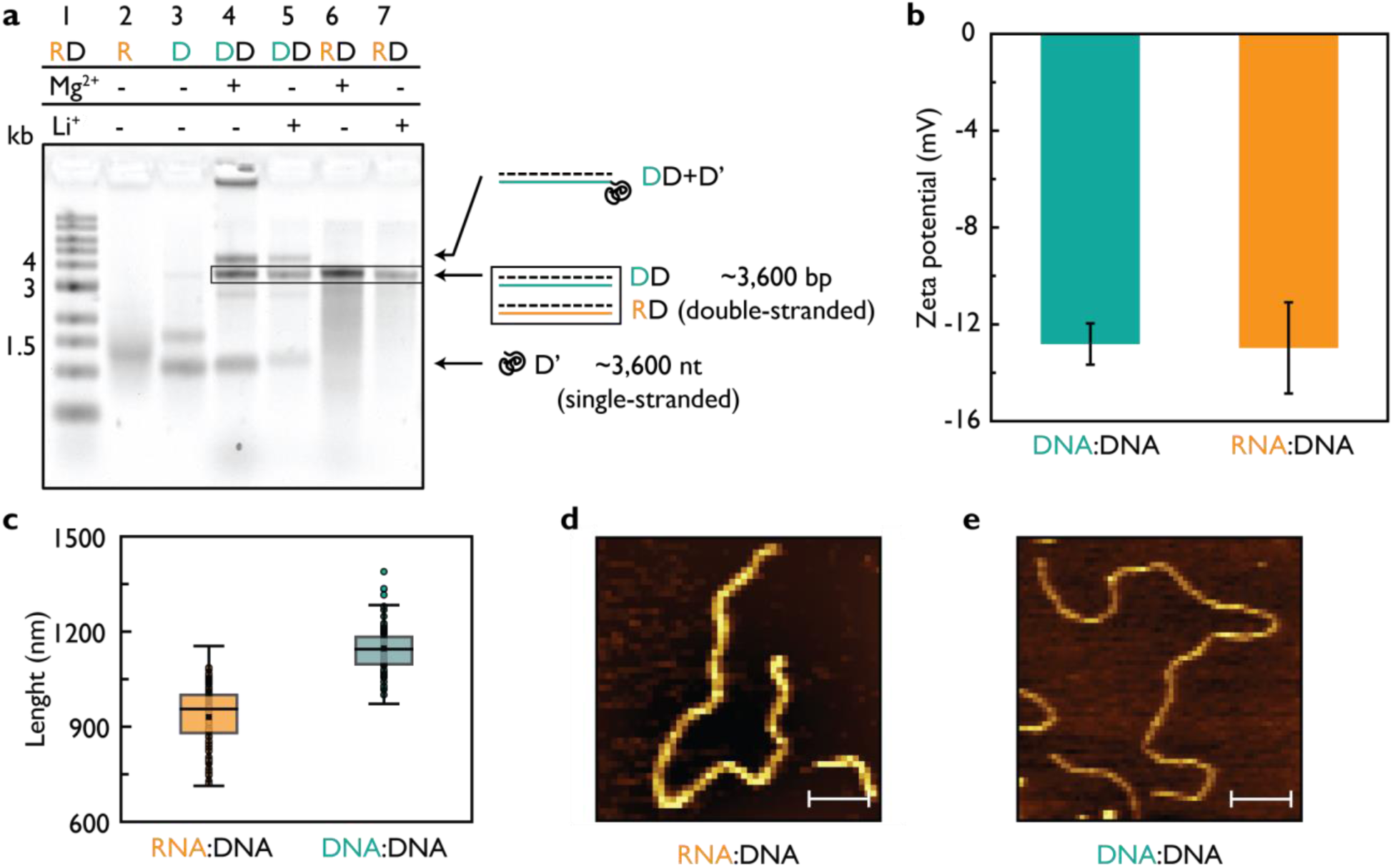
Length and charge of RD and DD were verified using agarose gel electrophoresis, DLS, and AFM imaging. (**a**) RD and DD duplexes were analyzed using 0.8% (w/v) agarose gel in 1 × TBE indicating a similar running speed. The gel lanes were as follows: 1 – 1 kbp ladder; 2 – MS2 RNA (3.6 kbp); 3 – cut ssM13 (7.2 kbp); 4 – assembled DD with ID ‘11001’ in 10 mM MgCl_2_; 5 – assembled DD with ID ‘11001’ in 100 mM LiCl; 6 – assembled RD ID ‘11100’ in 10 mM MgCl_2_; 7 – assembled RD with ID ‘11100’ in 100 mM LiCl; D’ represents to the half of ssM13 7.2 kbp (3,600 nt) that was not used for DD assembly. (**b**) The zeta potential of 26 bp DD (turquoise) and RD (orange) was measured in the nanopore measurement buffer (4 M LiCl, 1 × TE, pH 9.4) and found to be similar, both close to -13 mV. The sample size consisted of three technical replicates with 30 runs. (**c**) AFM imaging of ≈3.6 kbp RD and DD revealed contour lengths of 906.5 ± 8.7 nm and 1144.4 ± 5.8 nm (standard error of the mean), respectively. The sample size for RD and DD was N = 155 and N = 87, respectively. (**d**) An example AFM image illustrating the RD duplex. (**e**) An example AFM image illustrating the DD duplex. The scale bars for (**d**) and (**e**) are 100 nm.

The zeta potential, which is an important parameter often used to predict nucleic acid duplex velocity, was determined for RD and DD duplexes of the identical base pair length (26 bp) using DLS (**Figure 3b**). The zeta potential measurements demonstrated that RD and DD possess similar zeta potential values, indicating no significant difference in their surface charge. The theoretical surface charge of RD and DD differs by approximately 10% if calculated based on the surface charge of a cylinder using the equation *σ* = *q*/2*πrh*, where *σ* represents the surface charge, *q* = 2*e* is the charge per base pair, *r* is the duplex radius and *h* is the base pair rise. Hence, the measured zeta potentials of RD and DD align within the margin of error, confirming the agreement of their surface charges.

To validate the contour length of RD and DD, we performed AFM imaging (**Figure 3c-e; Figure S11**). We measured the length of ≈3.6 kbp RD and DD molecules from AFM images using Fiji ImageJ. ^39^ The measured length of RD was found to be 906.5 ± 8.7 nm, while for DD it was 1144.4 ± 5.8 nm (**Figure 3c**). These values for RD and DD correspond to base pair rise of 2.54 Å and 3.2 Å, respectively. These measurements are consistent with the values previously obtained from AFM imaging. ^40^ The AFM images of RD molecules are shown in Figure S11a, with an example displayed in **Figure 3d**. These images predominantly show duplex structures. On the other hand, AFM images of DD (Figure S11b; **Figure 3e**) exhibit multiple species, as observed in the agarose gel analysis (**Figure 3a**), including coiled single-stranded D’ and DD + D’ formed from single-cut M13. The AFM images also reveal the presence of single-stranded coils originating from non-complemented ssM13 DNA (Figure S11b). The AFM results regarding the length of the molecules support the assumption that RD adopts an A-form-like duplex conformation, while DD adopts a B-form duplex conformation.

### Molecular dynamics simulations elucidate the force per unit length exerted on RD and DD duplexes

We performed all-atom molecular dynamics (MD) simulations to characterize the effective electrophoretic force acting on RD and DD constructs. A typical simulation system contained a nucleic acid duplex placed in a periodic box of solvent, **Figure 4a**. In a force measurement simulation, a 25 mV/nm electric field was applied along the duplex while a harmonic restraint opposed the displacement of the center of mass of the duplex. The resulting effective force, which is equal by magnitude and opposite in direction to the force of the restraint, was greater on DD than an RD duplex, **Figure 4b**, similar to that reported previously. ^41^ When the force is divided by the base pair rise of the respective duplex, the all-atom simulations yield 20.98 and 19.47 pN/nm for DD and RD duplexes, respectively, corresponding to an approximately 8% higher force per unit length on DD duplexes (**bar graph inset, Figure 4b**). The force exerted per unit length is similar between RD and DD and they have similar velocities. Hence, we would expect that A-form RR duplexes of the same length will have a similar velocity to the studied ones in our study.

**Figure 4.**
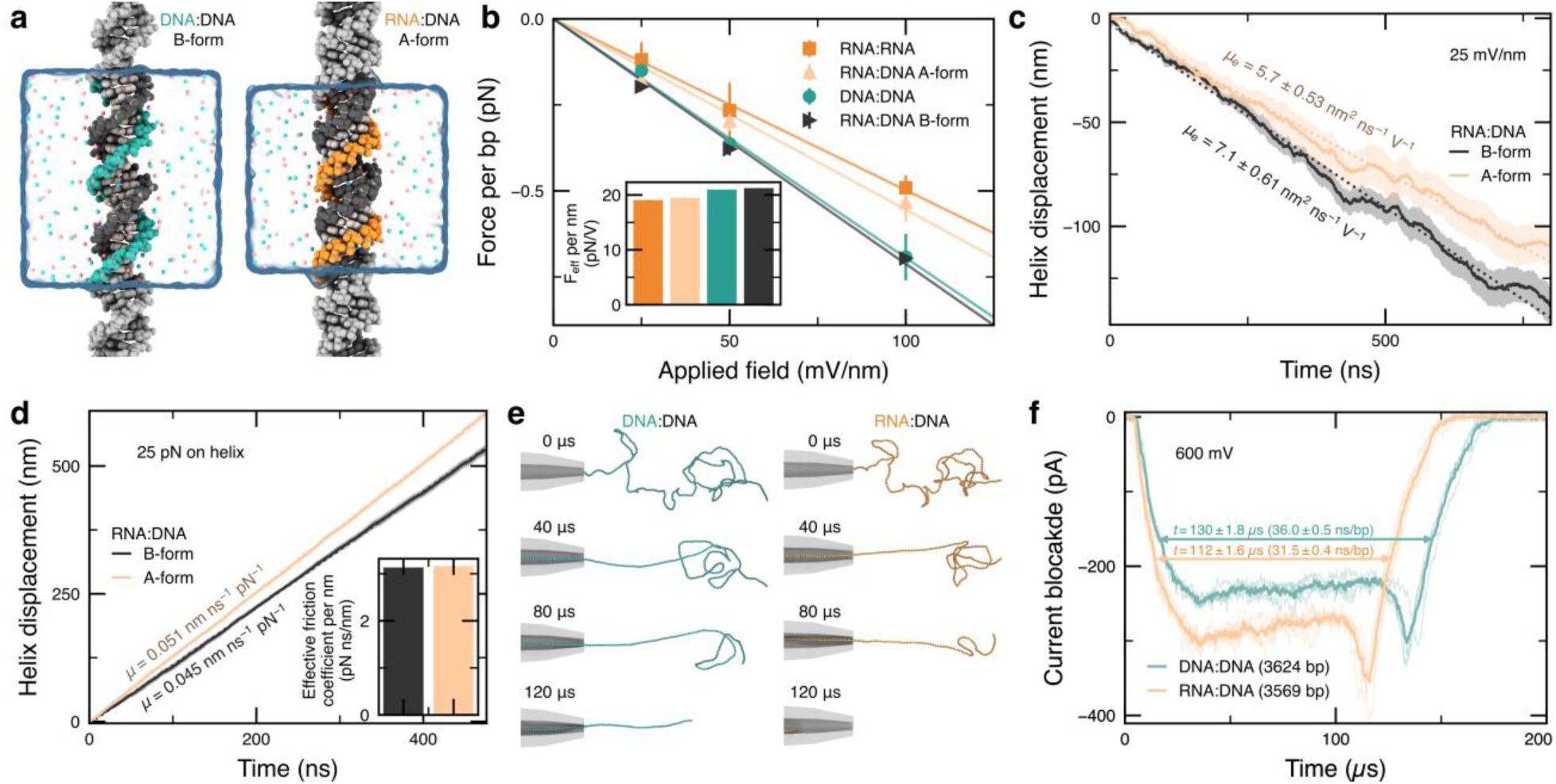
Simulations of DD and RD duplexes. (**a**) All-atom MD simulation systems, each containing two helical turns of nucleic acid duplex solvated in 4 M LiCl electrolyte (turquoise and orange spheres depicted in the 9.4-Å thick slab centered on the duplex). (**b**) Dependence of the effective force on duplexes obtained from a harmonic restraining potential that held the center of mass of the duplex at the origin while an electric field was applied in all-atom simulations. The inset depicts the effective force per unit length of duplex. (**c, d**) Displacement of duplexes in all-atom simulations under a 25 mV/nm applied field (**c**) or a 25 pN applied force (**d**). The slope of the displacement provides an estimate of the electrophoretic and hydrostatic mobilities, respectively. The inset in panel (d) depicts the dependence of the friction coefficient for a unit length of duplex estimated from the hydrostatic mobility. (**e**) Snapshots depicting the translocation of 3.6 kbp DD and RD duplexes through a conical nanopore in coarse-grained BD simulations. The mobility and effective electric force acting on the duplexes was taken from the all-atom MD simulation data. (**f**) Nanopore ionic current blockades estimated from five (RD) and four (DD) replicates of the DD and RD duplex translocation simulations. The bold lines depict the average current among the replicas.

When an RD duplex was constrained to a B-form configuration, the observed force matched exactly the force of a DD duplex (**Figure 4b**), indicating that the A-form geometry reduces the force per base pair rather than the additional hydroxyl group in the RNA. Additional simulations of finite RD duplexes not bound across the periodic boundary starting in A-form-like and B-form geometries revealed that B-form RD duplexes are unstable, while A-form-like RD duplexes are stable in 4 M LiCl, **Figure S12a**. Consistent with the lower effective electric force, A-form-like RD duplexes were observed to translate more slowly relative to the surrounding solvent compared to B-form RD duplexes in simulations of periodic duplexes under applied field and no restraints, **Figure 4c**. However, when a 25 pN force was applied directly to the helix, an A-form-like RD duplex was seen to move faster than an unstable B-form RD duplex, **Figure 4d**. Finally, the motions of the ions were analyzed to determine a dependence of the conductivity of 4 M LiCl solution on the distance from a RD or DD duplex, **Figure S12d**.

The mobility results from the all-atom MD simulations were incorporated into a coarse-grained Brownian dynamics (BD) model of long duplexes undergoing nanopore translocation, **Figure 4e**. The duplex models used two beads to represent every base pair with the beads connected by harmonic potential that recapitulate the experimental persistence lengths (DD: 50 nm; RD: 60 nm) using data for RR as a proxy for RD duplexes. The current blockade was estimated during BD simulations of translocation for each 3.6 kbp DD and RD duplex, **Figure 4f**. The simulated ionic current traces were found to reproduce the overall shape of the experimental current blockades (as shown in **Figure 2**) with the exception of a current dip at the end of the translocation events, which close inspection of the simulation trajectories attributed to DNA recoiling after its passage through the nanopipette aperture. In our simulations, the RD duplex is found to translocate faster, that is in qualitative agreement with the trend observed for nanopore translocations of 3.6 kbp RD and DD duplexes. In similar simulations performed with a longer pipette and a 2.7 kbp DD duplex of a contour length similar to that of the 3.6 kbp RD duplex, the RD duplex we find that the translocation times of 2.7 kbp DD and 3.6 kbp RD are within the error of our experimental nanopore measurements (**Figure S13**). The observed importance of the contour length is in agreement with the experimental results within experimental errors. Additional simulations performed using the 2.7 kbp DD and 3.6 kbp RD duplexes for a variety of pore geometries, including a nanoslit, ^42^ a cylindrical nanopore and an hourglass nanopore, revealed that the ratio of the DD to RD duplex translocation times does not depend on the pore geometry or shape (**Figure S14**).

### Theoretical basis of nanopore translocation velocity

Neglecting the possibility of direct interactions between a nucleic acid duplex and a nanopore surface, a line charge density, defined as the number of charges per base pair, can be used to estimate the force *F_xx_* on the nucleic acid molecules in nanopores as 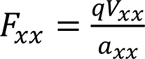, ^43^ where *q* is the charge per base pair, *V_xx_* the applied potential over the nanopore, and *a_xx_* the rise of RD or DD duplex. Setting *q* = 2*e* and assuming *V_RD_* = *V_DD_* in the same nanopore, we estimate the force ratio of RD over DD as

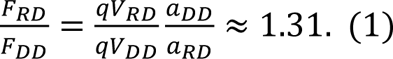

The result of ∼1.3 greatly overestimates the force acting on the A form and contradicts both the experimental results as well as the simulations.

A more accurate view of the forces determining translocation times is achieved by considering the hydrodynamic friction with the solvent. Considered our recent results on the time dependence of translocation time on molecular configuration, ^44^ we assume the limiting case that both RD and DD molecules are completely stretched out cylinders. The force ratio can then be calculated by assuming that the drag force is equal and opposite to the driving force. Using the experimentally obtained length of translocation time, τ and length, *L* for our RD and DD measurements we find (**Table S6**; the step-by-step calculations are shown in the **Section 1** and **2** of the Supporting Information):

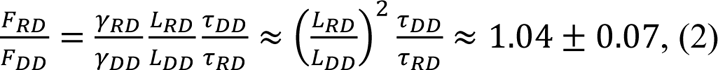

where γ is the approximated friction coefficient of a long, thin cylinder; *L* is the contour length of the duplex; and τ is the nanopore translocation time of the corresponding duplex, RD or DD. The friction coefficient for the limiting case of long rigid rods is 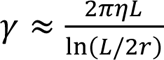 for both DD and RD. ^45^ Hence, the force ratio yields a value close to one using the experimental results. This is in excellent agreement with the MD simulations that revealed a force ratio per contour length differs by a few percent. Both AFM imaging and MD simulations demonstrated that the RD duplex indeed is closer to A-form while the DD duplex has B-form. However, when we tried to force the RD duplex to be B-form, it was unstable in the MD simulations, further confirming that the contour length is as postulated. The zeta potential, an indicator of the surface charge, is also similar for both the RD and DD duplex.

## DISCUSSION

In this study, we have determined that the time a nucleic acid duplex takes to translocate through a nanopore is primarily determined by the duplex’s contour length. Through experiments involving duplexes of identical base pair numbers but adopting different structural conformations—A-form-like and the canonical B-form—we observed that the A-form-like requires less time to translocate through the nanopore. We attribute this observation to the differences in the contour length.

Our experiments performed using RD duplexes, which have a larger persistence length than DD duplexes, and experiments performed using DD duplexes that contained or lacked nicks in one of the DNA strands, suggest that neither persistence length of the duplex nor the presence of the nicks affect the translocation time under our experimental conditions. Besides, there are other extreme scenarios regarding the influence of the nanopore diameter on the duplex translocation. ^10,15,38,46–49^ That is, when the pore diameter is comparable to the diameter of the duplex, the duplex-pore interactions become pivotal. Here, the direct contact force between the duplex and the nanopore wall becomes important. ^47^ Conversely, when the pore diameter surpasses the duplex’s persistence length, ^49,50^ the persistence length might have more profound effects on the translocation time. Specifically, the force exerted on the duplex outside the nanopore has significant effects on the translocation time. We note that, according to Equation 1, the effective force per unit length may depend on the duplex diameter for duplexes shorter than 50 nm.

When discussing the electrophoretic transport of nucleic acids, reliance solely on gel electrophoresis data presents inherent challenges. ^51,52^ Notably, the correlation between gel type and percentage with the translocation behaviors of RD- and DD-duplexes is not always linear, with interactions between the gel matrix and the duplexes significantly influencing the outcomes. ^51,52^ However, it is imperative to understand that gel characterization, while useful for determining relative DNA or RNA lengths, is not adept at discerning the relative translocation velocities of their movement within an electric field. Therefore, to address this intricacy, the adoption of nanopores combined with simulations emerges as a more precise and insightful approach.

## CONCLUSIONS

We have demonstrated that nanopore transport of double-stranded nucleic acids is influenced by the molecular structure of the duplex molecules, with RD (RNA:DNA) duplexes translocating faster than their DNA-only counterparts (DD).

Both experiments and simulations show that the force of an external electric field per unit length of RD- or DD-duplexes is similar within uncertainty of the measurement and, hence, the contour length of the molecule is a primary determinant of the translocation time.

Further studies involving different forms of nucleic acids such as RR duplexes, C-DNA, and Z-DNA in ionic liquids would contribute to a better understanding of molecular transport of nucleic acids ^53^. The higher velocity of RD duplexes has implications for the density of labels in RNA identifiers used for nanopore RNA analysis, ^32,54^ which is important for their design and detection. This poses a challenge when applying nanopores for the analysis of shorter RNA molecules (<2 kbp), and additional strategies for slowing down RNA identifiers may be necessary. Our study provides insights into the differences in molecular transport between non-canonical RD and DD nucleic acids, offering a proof-of-concept model of how charge and contour length influence translocation dynamics in electrophoretic transport. The deeper understanding of translocation dynamics of nucleic acids through nanopores is essential for accurate and precise analysis of RNA structural isoforms and RNA-molecule interactions.

## METHODS

### DNA scaffold preparation

In order to generate linear fragments of specific lengths, we performed a double-digestion on M13mp18 single-stranded circular M13 DNA (Guild Biosciences, 100 nM, 7,249 nt). In the first double-digestion reaction two DNA fragments of 3,621 nt (D) and 3,628 nt (D’) were generated using DraIII and BaeGI enzymes (New England Biolabs; **Figure S1a**). The digestion reaction involved annealing two oligonucleotides (**Figure S1b**) to the 7,249 nt circular scaffold, creating restriction sites for DraIII-HF and BaeGI-HF enzymes (**Figure S1c**). We mixed 8 µL of ssM13 scaffold with 1 µL of oligonucleotide A (100 µM, Integrated DNA Technologies), 1 µL of oligonucleotide B (100 µM, IDT), and in 1 × CutSmart buffer (NEB). The reaction mixture was pipetted ten times to ensure thorough mixing, briefly spun down, heated to 70 °C for 30 s, and slowly cooled down to room temperature (20 °C) over 40 min. The resulting annealed structure was then combined with 1 µL of DraIII-HF (NEB; 100,000 units/mL) and 1 µL of BaeGI-HF (NEB; 100,000 units/mL) and incubated for 1 h at 37 °C. Subsequently, the mixture was purified using the Monarch PCR and plasmid DNA purification kit (NEB), eluted in 10 mM Tris-HCl pH 8.0 and stored at -20 °C for further experiments. The concentration of the cut scaffold was estimated using a NanoDrop UV-Vis spectrophotometer.

To prepare a DNA scaffold measuring 2,724 nt in length, we followed an identical protocol, but employed restriction endonucleases DrdI (NEB; 10,000 units/mL) and AfeI (NEB; 10,000 units/mL) instead, along with oligonucleotides C and D (**Figure S1 d-f**).

### Preparation of DNA:DNA identifier

To prepare the DNA:DNA identifier, we mixed 12 µL of the cut scaffold (30 nM) with 2.4 µL of complementary oligonucleotides specific to the desired fragment either for the 3.6 kbp or 2.7 kbp cut DNA scaffold (concentration of each oligo 1 µM, IDT). Additionally, we added 4 µL of a filtered solution of MgCl_2_ (100 mM, pH 7.4) or LiCl (1 M, pH 7.57), 4 µL of a filtered solution of Tris-HCl (100 mM, pH 8.0), and nuclease-free water to reach a total volume of 40 µL. The reaction was incubated at 70 °C for 5 min and then slowly cooled down over the course of 1 h to reach room temperature (20 °C). To remove excess oligonucleotides, the mixture was subjected to filtration using 0.5 mL Amicon Ultra filters with a 100 kDa cutoff. Specifically, the 40 µL reaction was combined with 460 µL of washing buffer (10 mM Tris-HCl, pH 8.0; 0.5 mM MgCl_2_), and centrifuged for 10 min at 9,300 × g at 4 °C. The flowthrough was discarded, and the previous washing step was repeated. After the second washing step, the filter was inverted and placed into a fresh 2 mL microtube. The filter was spun down for 2 min at 1,000 × g and 4 °C. The resulting sample was transferred to a 0.5 mL low binding DNA tube (Eppendorf) and stored at 4 °C until further use.

### Preparation of RNA:DNA identifier

To prepare the RNA:DNA identifier we combined 12 µL of 3,569 nt long MS2 RNA (Roche; 30 nM) with 2.4 µL of complementary oligonucleotides each at a concentration of 1 µM (IDT), 4 µL of a filtered solution of MgCl_2_ (100 mM, pH 7.4) or LiCl (1 M, pH 7.57), 2.9 µL of a filtered solution of Tris-HCl (100 mM, pH 8.0), and nuclease-free water to reach a final volume of 40 µL. The reaction was incubated at 70 °C for 5 minutes and then slowly cooled down over the course of 1 hour to reach room temperature (20 °C). Subsequently, the mixture was stored at 4 °C for future use. The preparation in MgCl_2_ is performed only for the assessment of the duplex behavior using agarose gel electrophoresis.

### Annealing of DNA and RNA duplexes

For the annealing process, we mixed the DD and RD 26 bp duplexes by combining oligonucleotides with the same sequence but different nucleic acid types (RNA or DNA). The reaction was performed in a solution containing 10 mM Tris-HCl (pH 8.0) and 100 mM LiCl (pH 7.57). The reaction was incubated at 70 °C for 5 minutes and then slowly cooled down over the course of 1 hour to reach room temperature (20 °C). Afterwards, the annealed duplexes were stored at 4 °C.

### Dynamic Light Scattering (DLS) of duplexes

To obtain the zeta potentials of the 26 bp DD and RD duplexes, measurements were conducted using a Malvern Zetasizer Nano ZSP instrument. The duplexes were prepared in nanopore measurement buffer with 4 M LiCl and 1 × TE pH 9.4. The DLS measurements consisted of 30 runs performed in triplicates to ensure accuracy and reliability of the data.

### Atomic Force Microscopy (AFM) imaging of identifiers

Atomic Force Microscopy (AFM) imaging of the 3.6 kbp long RD and DD duplexes was performed using an MFP-3D AFM System from Asylum/Oxford Instruments. The imaging was conducted in air using the non-contact mode. The duplexes were diluted to a concentration of 1 ng/μL in 1 mM MgCl_2_, and 10 μL of the solution was added to freshly cleaved mica. After a 1-minute incubation, the mica surface was rinsed with filtered Milli-Q H2O and then dried with nitrogen. The mica plate was affixed to the AFM sample stage using double-sided adhesive tape prior to scanning. Image visualization and analysis were carried out using Gwyddion software.

### Native agarose gel electrophoresis

To perform native agarose gel electrophoresis, we prepared a 0.8% (w/v) agarose gel in 1 × Tris/Borate/EDTA buffer (TBE) using RNase-free water. To inactivate potential RNases, we added 0.05% (v/v) bleach (NaOCl) to both the gel and running buffer. ^55^ The samples were mixed with 6 × purple loading dye without sodium dodecyl sulfate (NEB) and with 10 × TBE to make a 1 × solution. The gel was run in an ice bath for 2-3 hours at a constant voltage of 70 V using a BioRad electrophoresis power supply. After electrophoresis, the gel was washed with nuclease-free water and incubated for 10-15 minutes in 3 × GelRed (Biotium) staining solution. The post-stained gel was imaged using a UV lamp (Agitium), and the images were uniformly adjusted using the Fiji ImageJ plugin. ^39^ The adjustments included inverting the grayscale, subtracting the background using a 150-pixel rolling ball algorithm, and adjusting the brightness and contrast. A 1 kbp ladder (NEB) was used for the relative comparison of the bands.

### Nanopore measurements and data analysis

Nanopore measurements were recorded using an Axopatch 200B current amplifier and filtered at 100 kHz with a 1 MHz sampling rate. All the measurements were run in 4 M LiCl, 1 × TE, pH 9.4 under 600 mV applied voltage. We prepared the duplexes with monovalent streptavidin in 0.5 mL DNA low binding tubes (Eppendorf). We pipetted components in this order: 4 M LiCl, 1 × TE pH 9.4, then the 8 M LiCl volume equal to the volume of the assembled RD and DD duplexes and monovalent streptavidin, then monovalent streptavidin in excess, and finally the assembled DD and RD duplexes to 0.3-0.5 nM. We mixed the sample by pipetting ten times in and out and loaded it on the nanopore chip. The molar ratio of single-stranded scaffold (MS2 RNA or M13 DNA): biotinylated overhangs: monovalent streptavidin was 1:3:9 (up to 1:3:30). Custom-built LabVIEW codes ^37^ were utilized for data recording and analysis. In brief, individual nanopore events were separated based on the minimal current drop threshold (80 pA), minimum duration (0.1 ms), and a manually adjusted range of event charge deficit (ECD) that typically fell within the range of 10 to 500 fC. These isolated nanopore events were further selected to exclude folded molecules, fragments, and aggregates. ^32,56^ The duplexes can fold anywhere along the molecule, ^24,57^ which limits their classification in their folded conformation. That is the reason why we only employ unfolded, linear duplex nanopore events. Hence, only the unfolded, linear RD and DD nanopore events were used for identification.

### All-atom MD simulations of short RD and DD duplexes

Except where specified, all-atom MD simulations were performed using the NAMD package, ^58^ the CHARMM36 additive force field ^59,60^ with CUFIX corrections for interactions between ions and phosphates, ^61^ periodic boundary conditions, smooth truncation of short-ranged non-bonded atomic interactions between 8 and 10 Å, and particle mesh Ewald summation ^62^ with a 1 Å grid spacing for long-range electrostatics. Hydrogen mass repartitioning ^63^ enabled use of a 4-fs timestep, while hydrogen bonds were constrained by SHAKE and RATTLE algorithms. ^64^ A Langevin thermostat applied to non-hydrogen atoms held the temperature constant at 291 K with a damping coefficient of 0.1 ps^−1^. In constant pressure simulations, a Langevin piston barostat was employed to maintain a pressure of 1 atm with piston period and decay times of 2 and 1 ps, respectively, with the cell basis vector along the helical axis fluctuating independently from the other axes. Where specified, the colvars module ^65^ of NAMD was used to restrain the center of mass of all phosphorus atoms using a spring constant of 500 kcal mol^−1^ Å^−2^. Atomic coordinates were recorded every 25,000 steps.

The simulation systems were assembled using idealized configurations for the duplexes (B-form: 21 bp and 3.4 Å rise; A-form: 22 bp and 2.6 Å rise) before solvating with a pre-equilibrated patch of TIP3P water. Water molecules were randomly replaced with ions to neutralize the system and provide a 4 M LiCl solution. The configurational energy of each system was minimized through 1000 steps of the conjugate gradient method, followed by 3-4 ns of simulation with non-hydrogen duplex atoms harmonically restrained about their initial positions (*k*_spring_ = 0.2 kcal mol^−1^) and the volume held fixed, followed by at least 5 ns equilibration with a barostat. In subsequent production simulations with an electric field or constant external force applied, the system volume was held constant. Four to eight replicas of each system were used for the production simulations with total sampling of at least 400 ns for each data point.

### Coarse-grained simulations of the full-length RD and DD duplexes

First, a continuum model of the glass nanopore (nanopipette) was constructed using COMSOL with *Electrostatics*, *Transport of Diluted Species*, and *Creeping Flow* physics modules used to solve an axisymmetric 2D model. The length of the pipette was set to 300 nm, the inner radius of the aperture to 5 nm, the pore widening angle to 4.87° for the first 50 nm and 1.2° for the rest of the pipette. The chambers of solution on either side of the pipette were 300 nm long (along the axis of the pore) and 150 nm in radius. The ion concentration was set to 4 M and the diffusion coefficients were set to 1.030×10^−9^ and 2.032**×**10^−9^ m^2^/s for Li^+^ and Cl^−^, respectively. The opposing ends of the two chambers provided 600 mV and electrostatic ground, zero pressure, and constant ion concentration boundary conditions. The walls of the chambers and the nanopore were subjected to zero ion flux and no-slip fluid boundary conditions. Zero and –0.01 C/m^2^ charge density was set as the boundaries of the chambers and at the surface of the pipette, respectively. The COMSOL modules were coupled *via* a volumetric charge density prescribed by the local concentration of ion species, the convective motion of the fluid acting on the ions, and a volume force acting on the fluid due to the ions. The resulting solution for the electrostatic potential was exported as previously described ^66^ using custom Python scripts to interpolate the points to a regular cartesian lattice with a 10 Å spacing along the pore axis and a 5 Å spacing in the orthogonal directions. A steric potential *u* was generated in regions where the COMSOL solution was invalid (*i.e.* inside the pore walls) by calculating the distance *d* to the nearest voxel with a valid solution, and setting the steric potential to *u* = *k_spring_* × *d*^2^, where *k_spring_* = 4 kcal mol^−1^ Å^2^. An algorithm was then used to iteratively fill invalid voxels of the electrostatic potential with the average value from immediately neighboring valid (or previously filled) voxels. The potential acting on each bare charge of a DNA duplex was reduced by a factor of 1/32 to represent the effective force. An identical protocol was used to construct potentials for the 1000 nm long conical pore described in Figure S13 and the cylindrical and hourglass shaped pores in Figure S14, except the pore profile and thickness of the membrane were updated. The nanoslit depicted in Figure S14 was constructed using a 3D COMSOL model as previously described, ^42^ except the ion conductances and voltages were set to match the conditions of the other pores in this study, and no surface charge was applied to the walls of the slit.

Using the multiresolution mrDNA Python package, ^67^ a model of a 2.7 kbp B-form DD duplex molecule was relaxed in three simulations with resolutions of 10 and 5 bp/bead and 1 bp/2 beads lasting 120, 6, and 2.4 μs respectively (timesteps of 200, 100, and 40 fs). During the simulations, a harmonic restraint (*k*_spring_ = 1 kcal mol^−1^ Å^2^) applied to one end of the DNA was moved from 25 nm outside to 12.5 nm inside the pore. The pore was represented only through the steric potential during the simulations. This process was performed eight times to provide unique initial conditions for subsequent production simulations that employed both electric and steric potentials, used a 1 bp / 2 bead model with 40 fs timestep, and lasted at least 300 μs until the duplex had translocated most of the way through the pore.

For the RD duplex, the same contour path at the end of the equilibration simulations was used to initialize the coordinates of a 1 bp / 2 bead model of a 3.6 kbp A-form duplex, adapted from mrDNA to have potentials consistent with an inter base pair spacing of 2.77 vs 3.4 Å, ^68^ a helical pitch of 11.6 vs 10.44 bp (32.1 vs 35.5 Å), an elastic constant of 560 vs 1000 pN, ^68^ and a persistence length of 60 versus 50 nm. The non-bonded interactions from the mrDNA package were unmodified for RD duplexes, and the width of the duplex was only incorporated implicitly through the modified diffusion coefficients and electrophoretic scaling factor used to convert potential to force, both quantities being extracted from the all-atom MD simulations as follows. For RD duplexes, the force due to the electrostatic potential was scaled by the ratio of the effective electrophoretic forces acting on non-canonical RD duplexes and B-form DD duplexes as observed in our all-atom MD simulations. Additionally, the diffusivity of each bead was scaled by the ratio of RD to DD hydrostatic mobilities as observed in all-atom MD simulations. Prior to production simulations, each RD system was relaxed with only the steric pore potential applied during an additional 2.4 μs simulation, during which the linking number of the duplex was allowed to relax.

After the completion of the trajectories, the steric exclusion model (SEM) ^69^ was used to calculate the ionic current blockade in each system as described previously, except the results of the ionic conductivity from all-atom MD simulations were used to compute the dependence of the conductivity of the solution on the distance from the duplex. ^70^ The experimental values for the bulk conductance near 4 M LiCl ^71^ was interpolated to provide the value of 151 mS/cm used in the SEM calculations.

## ACKNOWLEDGMENTS

F.B. acknowledges research funding from the George and Lilian Schiff Foundation Studentship, the Winton Programme for the Physics of Sustainability PhD Scholarship and St John’s College Benefactors’ Scholarship. G.P.G. acknowledges funding from EPSRC CDT MRes/PhD Studentship in Nanoscience and Nanotechnology (NanoDTC Cambridge EP/S022953/1) and Trinity-Henry Barlow Scholarship. R.T. and U.F.K. acknowledge funding from an European Research Council (ERC) consolidator grant (DesignerPores no. 647144), an ERC-2019-PoC grant (PoreDetect no. 899538), and UK Research and Innovation (UKRI) under the UK government’s Horizon Europe funding guarantee EP/X023311/1. R.T. acknowledges funding from the European Union’s Horizon 2020 research and innovation programme under the Marie Sklodowska-Curie Grant Agreement No. 892333, and from the Blavatnik Family Foundation. C.M., and A.A. acknowledge funding through the Human Frontier Science Project (RGP0047/2020) and the Leverhulme Visiting Professorship grant [VP2-2019-012] to AA. ARBD development is supported by the National Science Foundation grant OAC-2311550. The supercomputer time was provided through XSEDE Allocation Grant MCA05S028 and the Leadership Resource Allocation MCB20012 on Frontera of the Texas Advanced Computing Center.

