## Supplementary Information for "Nanopore translocation reveals electrophoretic force on non-canonical RNA:DNA double helix"

**This PDF file includes**

Materials and Methods
Supplementary Text
Figs. S1-S15
Tables S1-S7

**Supplementary Materials and Methods**

Materials

The commercial buffers used in this study were Tris-EDTA buffer solution 100 × concentrate (Sigma-Aldrich, catalog number T9285), 0.2 µm filtered 1M $MgCl_2$ (Invitrogen by Thermo Fisher Scientific, catalog number AM9530G), 0.2 µm filtered and autoclaved nuclease-free water (Ambion, catalog number AM9937). Lithium chloride for molecular biology ≥ 99% purity (Sigma-Aldrich, catalog number L9650), sodium chloride for molecular biology ≥ 99% purity (Sigma-Aldrich, catalog number S3014), Tris-HCl BioPerformance certified, ≥ 99% purity (Sigma-Aldrich, catalog number T5941). All buffers used in this study were filtered with 0.22 µm Millipore syringe filter units (Merck).

Glass quartz capillaries with filament (inner diameter 0.2 mm, outer diameter 0.5 mm) were purchased from Sutter Instrument Company. PDMS was purchased from Sylgard 184, Dow Corning (catalog number 101697), microscope slides clear ground 1.0 – 1.2 mm (Thermo Fisher Scientific, catalog number 1238-3118), silver wire with 1.0 mm diameter (Advent Research Materials Ltd, catalog number AG548711). Amicon 0.5 mL filter units (100 kDa cut-off) were purchased from Merck (catalog number UFC5100BK). Membrane Filter, 0.22 µm pore size membrane filters (MF-Merck Millipore™, catalog number GSWP04700).

DNA LoBind® Tubes (Eppendorf) were purchased from Thermo Fisher Scientific, and thin-walled, frosted lid, RNase-free PCR tubes (0.2 mL) were purchased from Thermo Fisher Scientific (catalog number AM12225).

RNA from bacteriophage MS2 3569 nt in length (Roche, catalog number 10165948001), total RNA from human cervical adenocarcinoma (Thermo Fisher Scientific, Invitrogen, catalog number AM7852) and human universal reference total RNA (Thermo Fisher Scientific, Invitrogen, catalog number QS0639).

Single-stranded circular m13mp18 7,249 nt in length (Guild Biosciences, foundation m13).

### Section 1: Electrophoretic forces acting on RD and DD in a nanopore are found to be similar

For double-stranded DNA the maximal driving force exerted on the molecule is often estimated using the line charge density $q/a$. The force is then given by $F = qV/a$, where $q = 2e$ is the charge per base pair, and $a = 0.34$ nm is the base pair distance [1].

If we calculate the ratio of forces exerted on RD and DD ($F_{RD}$ and $F_{DD}$, respectively) using the line charge density-based force equation $F = qV/a$, we obtain a value of $F_{RD}/F_{DD}$ around 1.3.

The Reynolds Number (Re) is a parameter used to determine the dominance of inertial forces in the translocation process through a nanopore. In the overdamped regime, where Re is low, inertial forces can be neglected [2] and polymer transport can be effectively described as flow through a pipe. We can estimate the magnitude $\mathrm{Re} = \rho u D_p/\mu$, where $\rho$ represents the density of the buffer solution, $u$ is the mean velocity of the fluid, $D_p$ is the nanopore diameter and $\mu$ is the dynamic viscosity of the buffer. In the case of solid-state nanopores, both the DNA velocity and the liquid velocity reach several mm/s. With $D_p \approx 10$ nm (**Figure S15**) and considering the density and viscosity of water, we find that $\mathrm{Re} = \frac{\rho u D_p}{\mu} \approx 10^{-5} \ll 1$. As expected, inertial forces are negligible and at the experimental timescales, all forces exerted on DNA sum to zero, $F_d + F_E = 0$ [2].

Here, we set the electric driving force as equal and opposite to the friction on the translocating molecule, $F_E = \gamma u$, where $\gamma(L, r)$ is the length and radius-dependent drag coefficient of the polymers studied, and $u$ is the velocity of the polymer chain. Previous investigations have examined the translocation process of unfolded DNA molecules through conical glass nanopores, revealing nearly constant velocity depending on the directionality of the translocation [3]. Besides, the driving and electrophoretic force in nanopores the electro-osmotic force originating from the charged nanopore surface is a significant factor [4]. In the simultaneous nanopore measurements of RD and DD, the electroosmotic force exerted on both molecules in the nanopore is assumed to be the same within experimental errors. For the subsequent analysis, a constant translocation velocity ($u$) will be assumed.

From the nanopore measurements, we obtained translocation times as in **Table S6** for 3.6 kbp DD and 3.6 kbp RD. In the low Reynolds Number regime, we can calculate the velocity of each construct as $u_{DD} = F_{DD}/\gamma_{DD}$ and $u_{RD} = F_{RD}/\gamma_{RD}$. The friction coefficient for the limiting case of long rigid rods is ($L \gg r$) as $\gamma \approx \frac{2\pi\eta L}{\ln(L/2r)}$ for both DD and RD [5]. In this limit, the radius of the constructs has a limiting contribution since it is under the logarithm, and we find $\frac{\gamma_{RD}}{\gamma_{DD}} \approx \frac{L_{RD}}{L_{DD}}$ as both constructs have the same number of base pairs. With the experimentally determined length of both constructs, we can relate the translocation times $\tau_{DD} = L_{DD}/u_{DD}$ and $\tau_{RD} = L_{RD}/u_{RD}$. Hence, we calculate the force ratio using nanopore translocation times as (Table S6):

$$(1)\ \frac{F_{RD}}{F_{DD}} = \frac{\gamma_{RD}}{\gamma_{DD}}\frac{L_{RD}}{L_{DD}}\frac{\tau_{DD}}{\tau_{RD}} \approx \left(\frac{L_{RD}}{L_{DD}}\right)^2 \frac{\tau_{DD}}{\tau_{RD}} \approx 1.04 \pm 0.07.$$

Our analysis suggests that the forces acting on both constructs have nearly the same value with $F_{RD}$ being larger than $F_{DD}$ by only a few percent. This is contrary to the line charge density-based force equation $F = qV/a$ which predicts a force ratio that is 30 % higher. The experimentally measured force ratio for RR and DD duplexes using optical tweezers is concordant with our data [1]. Gel electrophoresis measurements of both constructs indicate similar running speeds (**Figure 3a**). Zeta potential measurements of shorter constructs in the same buffer solutions show agreement within the experimental error (**Figure 3b**).

**Section 2: The force ratio calculations**

Using the equation 1:

$$(1)\ \frac{F_{RD}}{F_{DD}} = \frac{\gamma_{RD}}{\gamma_{DD}}\frac{L_{RD}}{L_{DD}}\frac{\tau_{DD}}{\tau_{RD}} \approx \left(\frac{L_{RD}}{L_{DD}}\right)^2 \frac{\tau_{DD}}{\tau_{RD}}$$

we can calculate the force ratio for different nanopore measurements.

In the first case, RD and DD were 3.6 kbp with their respective AFM lengths (base pair distance of 0.26 and 0.34 nm/bp, respectively) and with nanopore translocation times ($\tau$) (**Table S6**) we obtained the force ratio value of for seven different nanopore measurements:

$$\frac{F_{RD}}{F_{DD}} \approx \left(\frac{L_{RD}}{L_{DD}}\right)^2 \frac{\tau_{DD}}{\tau_{RD}} \approx 0.98 \pm 0.07$$

In the second case, RD and DD were 3.6 kbp and 2.7 kbp, respectively (base pair distance of 0.26 and 0.34 nm/bp, respectively) and with nanopore translocation times ($\tau$) (as shown in Figure 2d) we obtained the force ratio value of:

$$\frac{F_{RD}}{F_{DD}} \approx \left(\frac{L_{RD}}{L_{DD}}\right)^2 \frac{\tau_{DD}}{\tau_{RD}} \approx \left(\frac{3.6kbp * 0.26nm/bp}{2.7kbp * 0.34nm/bp}\right)^2 \frac{0.355 \pm 0.004ms}{0.351 \pm 0.004ms} \approx 1.05 \pm 0.01$$

**Section 3: The effects of nicks on the translocation velocity**

We constructed two DD structures (**Figure S10**). The first is the DD duplex `11001`, which is 3.6 kbp in length and contains 99 DNA nicks, as depicted in Figure 1 (using single-stranded circular M13 DNA (NEB, single-stranded circular M13mp18 DNA, N4040S) linearized as shown in Figure S1a,b; oligonucleotides are listed in Table S2). Subsequently, we created the same 3.6 kbp DD duplex from double-stranded M13 DNA (NEB, M13mp18 RF I DNA, N4018S) using an identical set of restriction enzymes yielding the DD duplex without DNA nicks. Both constructs were analyzed in the same nanopore measurement, as described in the Methods section.

The constructs were run on a 1 % (w/v) agarose gel, 1 × TBE (Figure S10a). We verified that 3.6 kbp DD constructs are made and then we proceeded to nanopore measurements

(Figure S10b-f; nanopore details are listed in Table S7). We calculated the percentage of unfolded (linear) and folded nanopore events (Figure 10b), showing no significant difference for two different nanopores. The translocation time and the scatter plot of the mean event current vs translocation time for unfolded events are displayed in Figure 10c and 10d, respectively. The higher mean event current for DD with nicks are expected due to the presence of the labels along the duplex.

**Figure S1.**

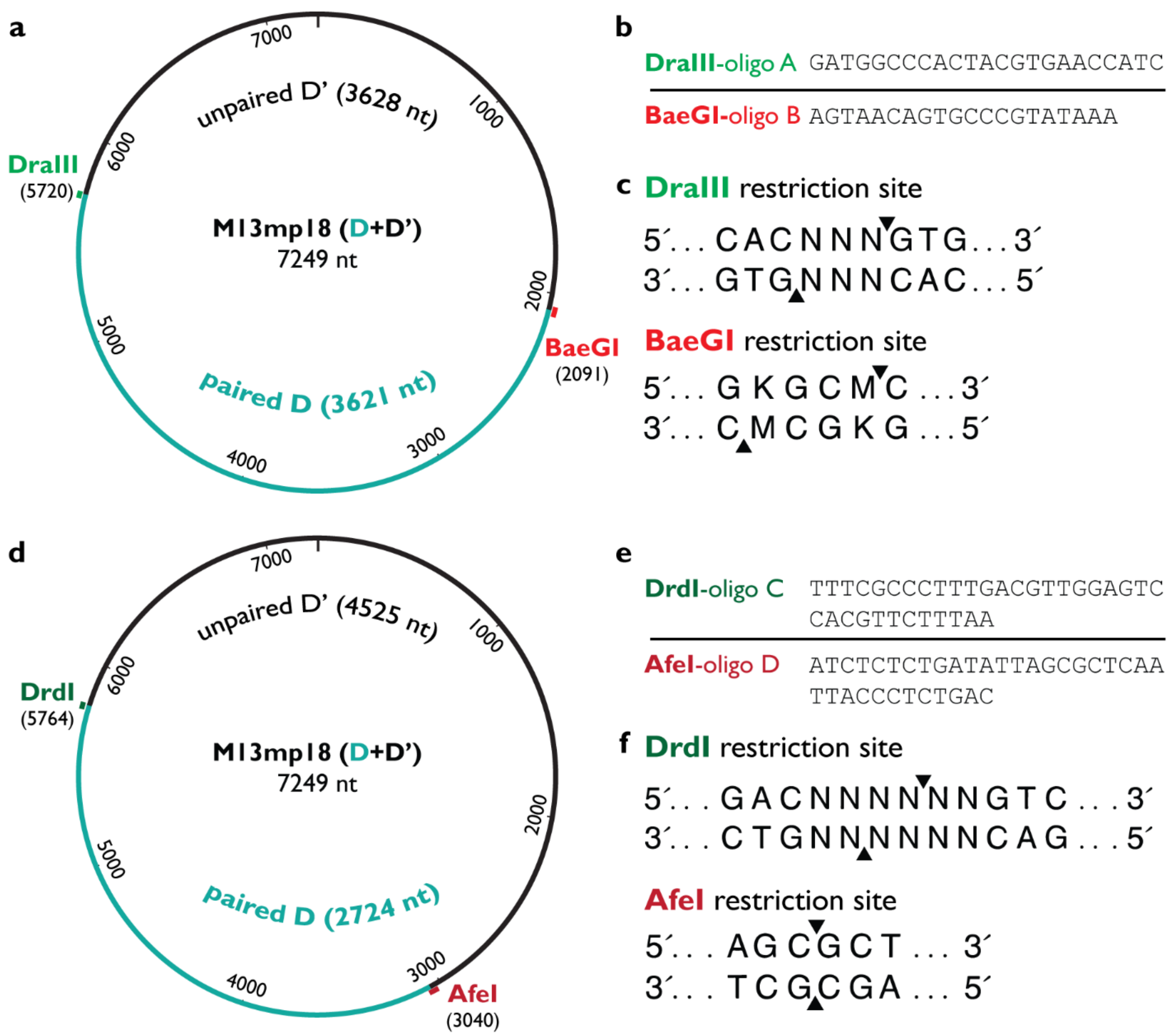


**Figure S1.** Annealing of oligonucleotides to single-stranded M13mp18 DNA (7,249 nt) to create cutting sites for producing single-stranded DNA fragments of desired length. (**a**) To create 3,621 nt long DNA fragment ssM13 was cut with DraIII and BaeGI restriction endonucleases. This double-digestion produces our desired fragment D (3,621 nt) that would be further paired with complementary oligonucleotides and the side fragment D' (3,628 nt). (**b**) Oligonucleotide sequences used for annealing to M13mp18 to make the cutting sites for DraIII and BaeGI. (**c**) Recognition sequences for DraIII and BaeGI with marked cutting positions in both strands are indicated. N – A, C, T or G; K – G or T; M – A or C. (**d**) To create 2,724 nt long DNA fragment ssM13 was cut with DrdI and AfeI restriction endonucleases. This double-digestion produces our desired fragment D (2,724 nt) that would be further paired with complementary oligonucleotides and the side fragment D' (4,525 nt). (**e**) Oligonucleotide sequences used for annealing to M13mp18 to make the cutting sites for

DrdI and AfeI. (**f**) Recognition sequences for DrdI and AfeI with marked cutting positions in both strands are indicated. N – A, C, T or G.

**Figure S2.**

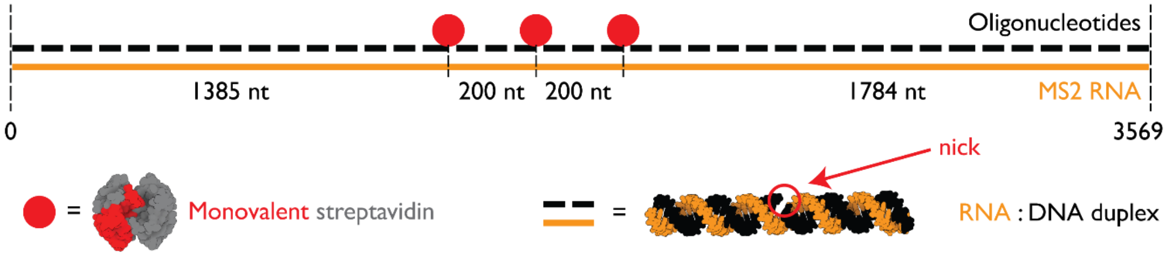


**Figure S2.** Detail design of 3.6 kbp RNA ID. The ID was assembled by binding short DNA strands (black; listed in Table S1) to 3,569 nt long MS2 phage RNA (orange). Monovalent streptavidins were bound to oligos with an overhang containing 3` biotins. Hence, creating a code `11100` where `1` represents single monovalent streptavidin bound to the 3`biotinylated oligonucleotide. These / sites are equally spaced by DNA strands. The whole RNA sequence has been fully base paired leaving only nicks. Nicks are discontinuities in only the DNA part of DNA:RNA molecule where there is no phosphodiester bond between adjacent nucleotides.

**Figure S3.**

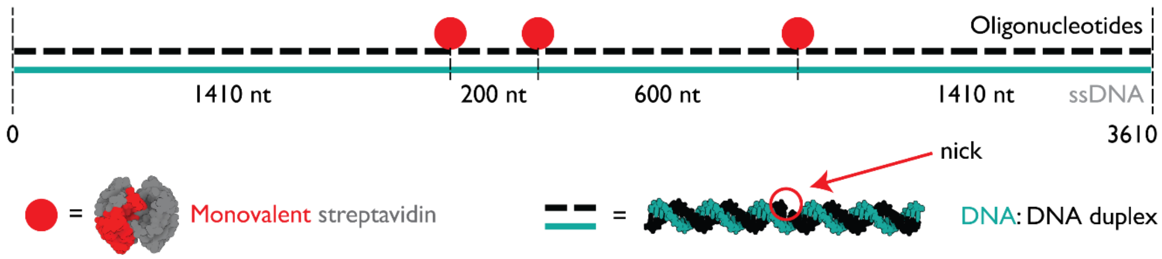


**Figure S3.** Detail design of 3.6 kbp DNA ID. The ID was assembled by binding short DNA strands (black; listed in Table S2) to 3,610 nt long double-digested m13mp18 phage DNA (turquoise). Monovalent streptavidins were bound to oligos with an overhang containing 3` biotins. Hence, creating a code `11001` where `1` represents single monovalent streptavidin bound to the 3`biotinylated oligonucleotide. These sites are spaced by DNA strands. The whole DNA sequence has been fully base paired leaving only nicks. Nicks are discontinuities in DNA:DNA molecule where there is no phosphodiester bond between adjacent nucleotides.

**Figure S4.**

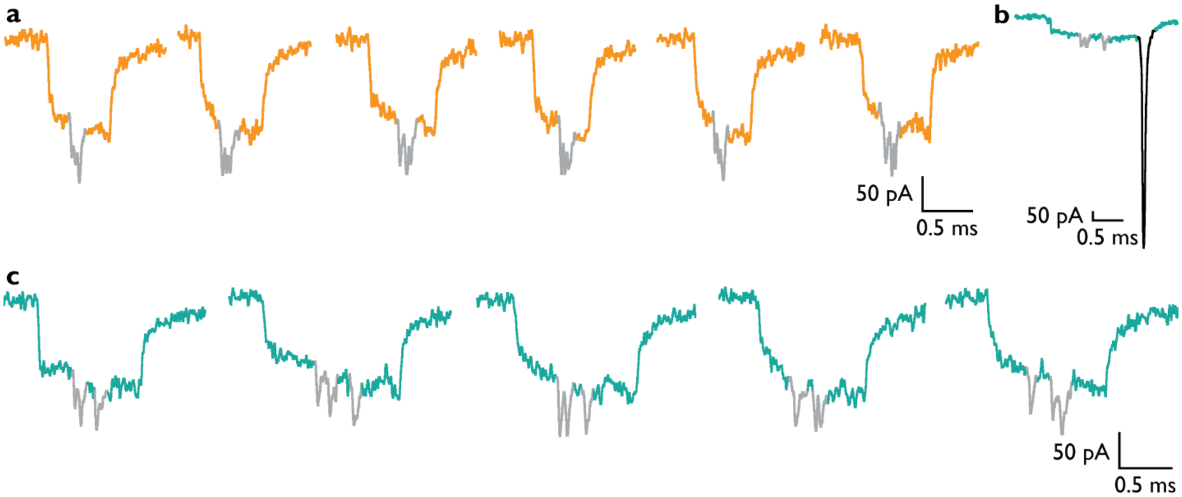


**Figure S4.** Nanopore example events for three species within a single measurement including (**a**) 3.6 kbp MS2 RNA ID, (**b**) 3.6 kbp M13 DNA ID with 3.6 kbp ssDNA part (in black), and (**c**) 3.6 kbp M13 DNA ID.

**Figure S5.**

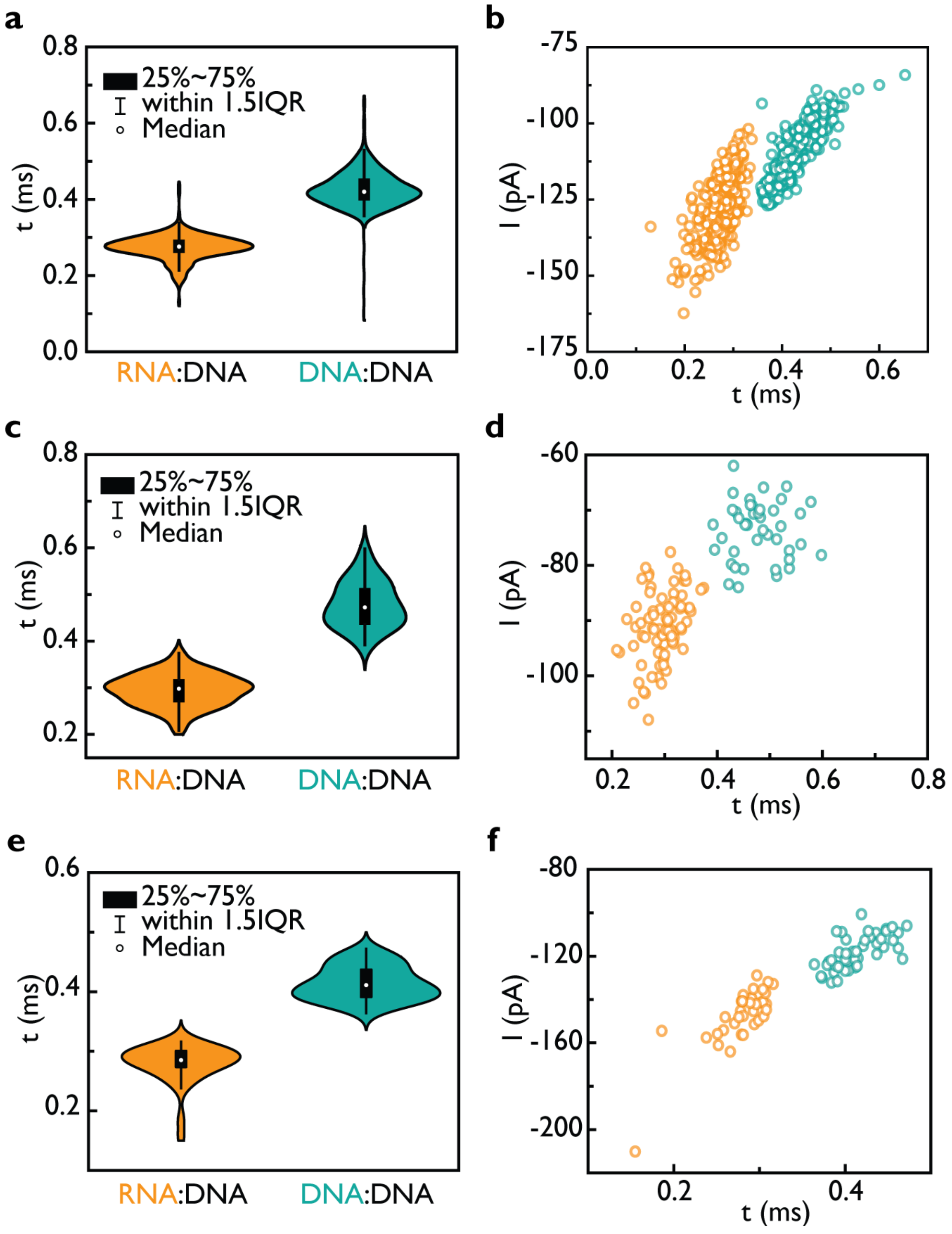
a
0.8
0.6
0.4
0.2
0.0
t (ms)
25%~75%
within 1.5IQR
Median
RNA:DNA
DNA:DNA
b
-75
-100
-125
-150
-175
I (pA)
0.0
0.2
0.4
0.6
t (ms)
c
0.8
0.6
0.4
0.2
t (ms)
25%~75%
within 1.5IQR
Median
RNA:DNA
DNA:DNA
d
-60
-80
-100
I (pA)
0.2
0.4
0.6
0.8
t (ms)
e
0.6
0.4
0.2
t (ms)
25%~75%
within 1.5IQR
Median
RNA:DNA
DNA:DNA
f
-80
-120
-160
-200
I (pA)
0.2
0.4
t (ms)

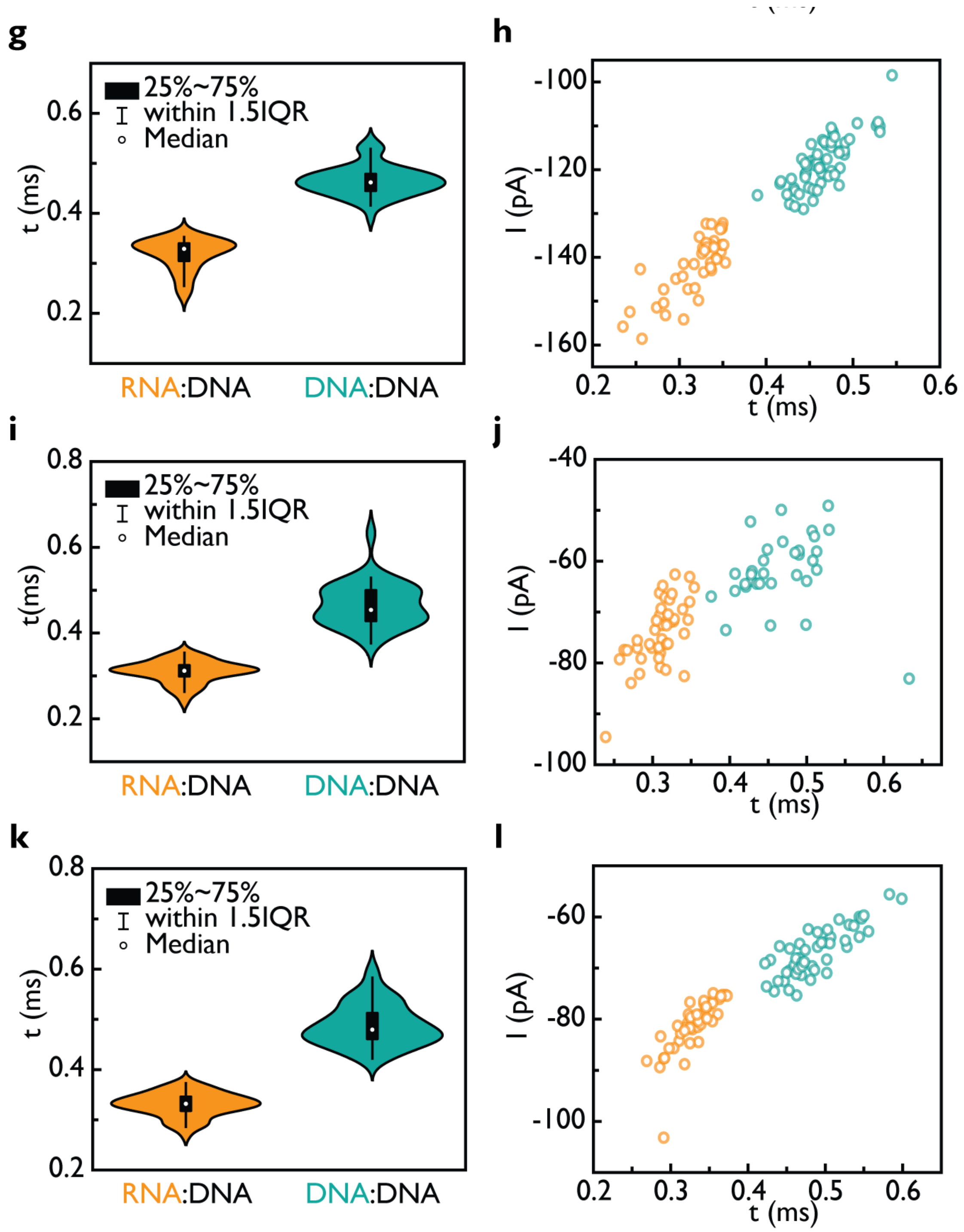


**Figure S5.** Nanopore data for 3.6 kbp RD and DD. Data for nanopore #2 are shown in (**a**) and (**b**). The sample size was 589. (**a**) The translocation time (τ) of RD and DD. (**b**) A scatter plot of the mean event current versus τ. Data for nanopore #3 are shown in (**c**) and (**d**). The sample size was 118. (**c**) The translocation time (τ) of RD and DD. (**d**) A scatter plot of the mean event current versus τ. Data for nanopore #4 are shown in (**e**) and (**f**). The sample size was 82. (**e**) The translocation time (τ) of RD and DD. (**f**) A scatter plot of the mean event current versus τ. Data for nanopore #5 are shown in (**g**) and (**h**). The sample size was 100. (**g**) The translocation time (τ) of RD and DD. (**h**) A scatter plot of the mean event current versus τ. Data for nanopore #6 are shown in (**i**) and (**j**). The sample size was 77. (**i**) The translocation time (τ) of RD and DD. (**j**) A scatter plot of the mean event current versus τ. Data for nanopore #7 are shown in (**k**) and (**l**). The sample size was 97. (**k**) The translocation time (τ) of RD and DD. (**l**) A scatter plot of the mean event current versus τ.

**Figure S6.**

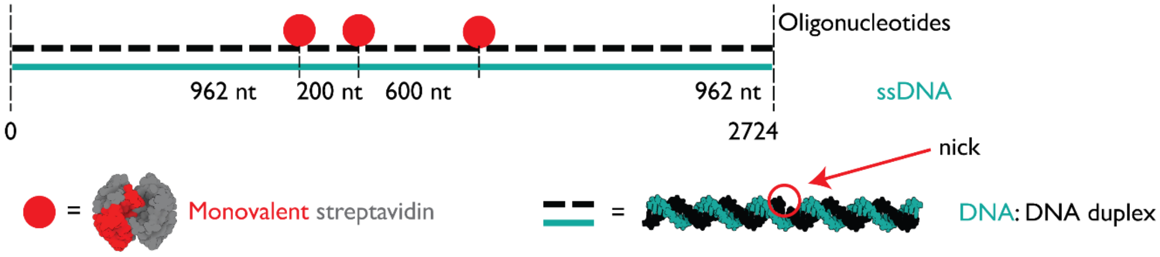


**Figure S6.** Detail design of 2.7 kbp DNA ID. The ID was assembled by binding short DNA strands (black; listed in Table S3) to 2,724 nt long double-digested m13mp18 phage DNA (turquoise).

**Figure S7.**

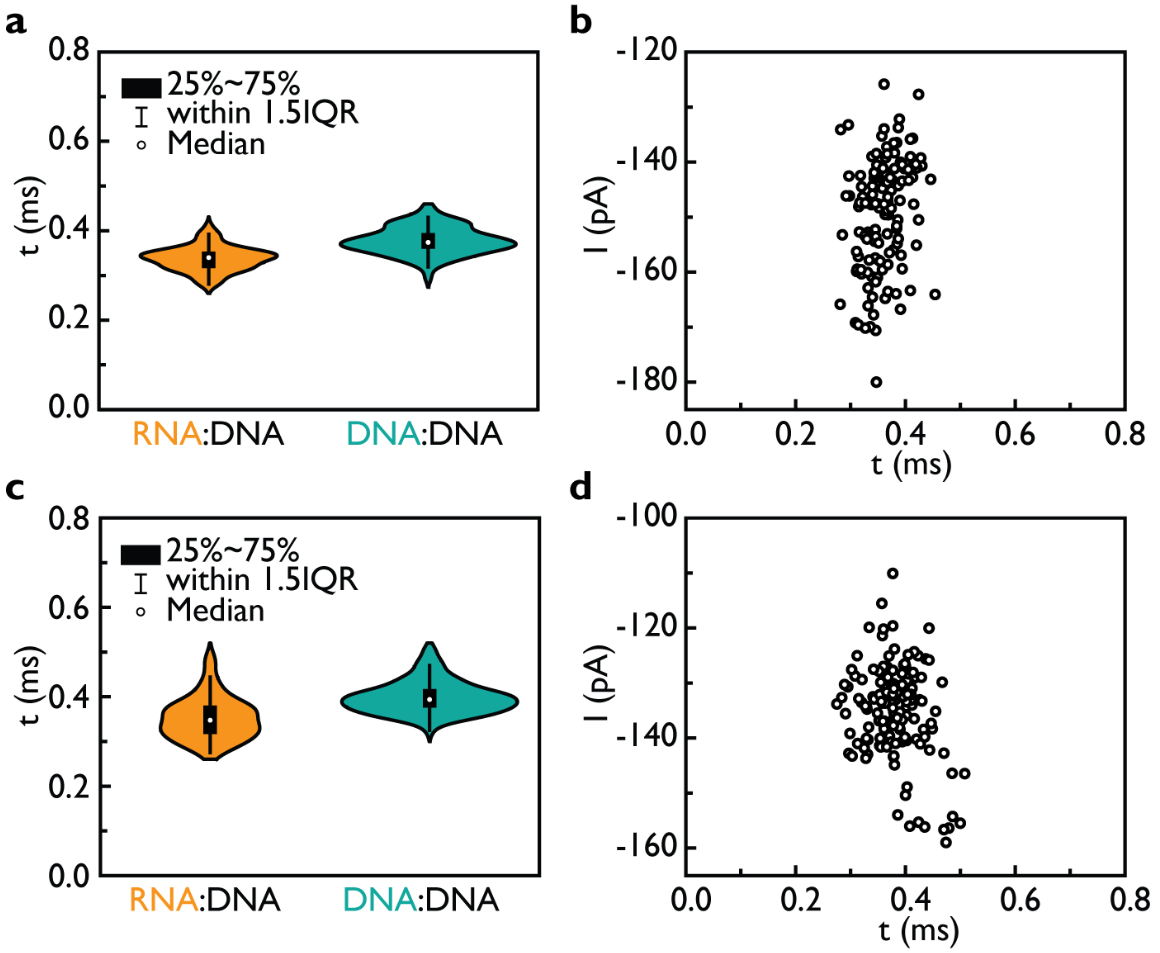
a
25%~75%
within 1.5IQR
Median
t (ms)
0.8
0.6
0.4
0.2
0.0
RNA:DNA
DNA:DNA
b
I (pA)
-120
-140
-160
-180
0.0
0.2
0.4
0.6
0.8
t (ms)
c
25%~75%
within 1.5IQR
Median
t (ms)
RNA:DNA
DNA:DNA
d
I (pA)
-100
-120
-140
-160
t (ms)

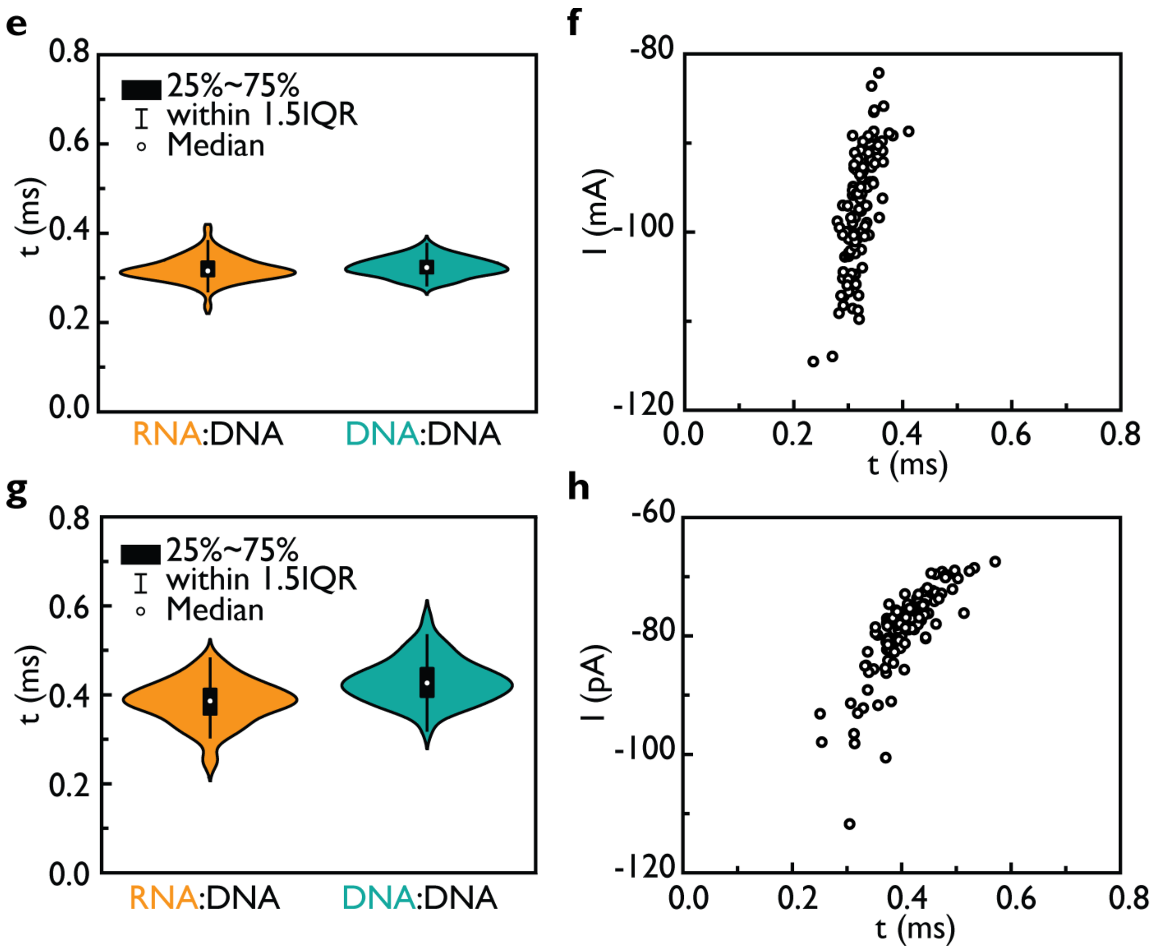


**Figure S7.** Nanopore data for 3.6 kbp RD and 2.7 kbp DD. Data for nanopore #2 are shown in (**a**) and (**b**). The sample size was 137. (**a**) The translocation time (τ) of RD and DD. (**b**) A scatter plot of the mean event current versus τ. Data for nanopore #3 are shown in (**c**) and (**d**). The sample size was 144. (**c**) The translocation time (τ) of RD and DD. (**d**) A scatter plot of the mean event current versus τ. Data for nanopore #4 are shown in (**e**) and (**f**). The sample size was 108. (**e**) The translocation time (τ) of RD and DD. (**f**) A scatter plot of the mean event current versus τ. Data for nanopore #5 are shown in (**g**) and (**h**). The sample size was 113. (**g**) The translocation time (τ) of RD and DD. (**h**) A scatter plot of the mean event current versus τ.

**Figure S8.**

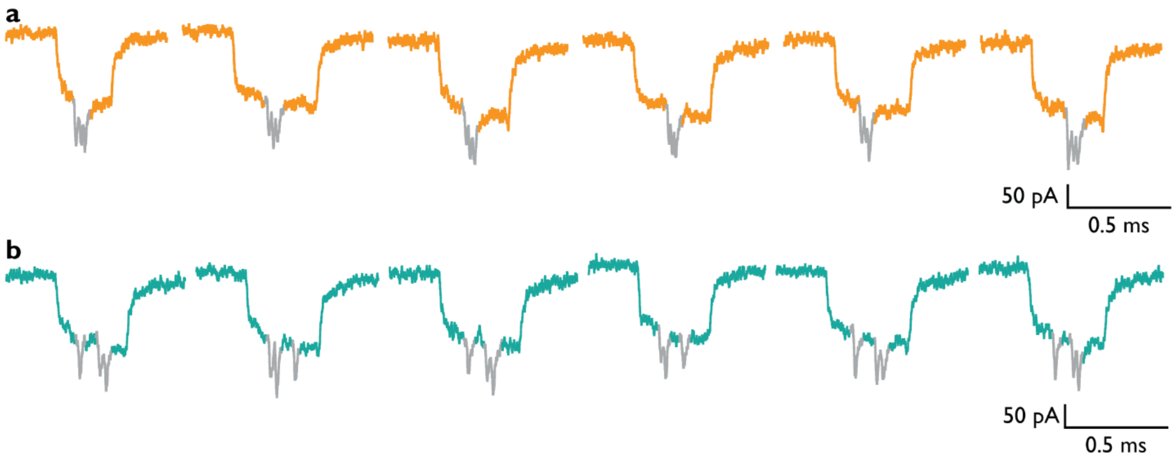


**Figure S8.** Nanopore example events for three species within a single measurement including (**a**) 3.6 kbp MS2 RNA ID, (**b**) 2.7 kbp M13 DNA.

**Figure S9.**

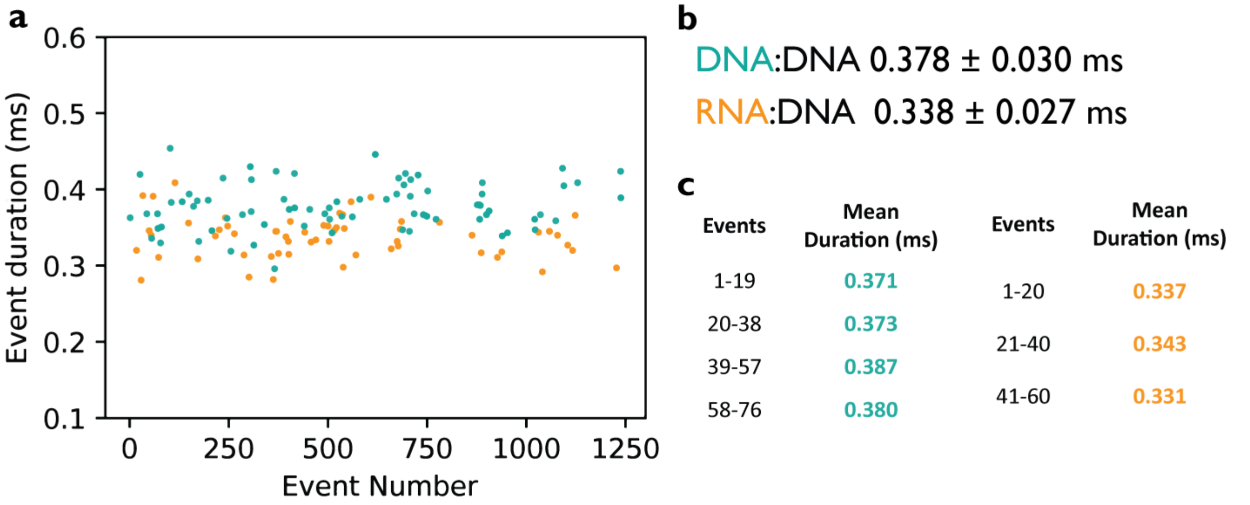


**Figure S9.** The unfolded 3.6 kbp RD and 2.7 kbp DD duplex translocation time with the event number does not alter significantly. (**a**) Average translocation time plotted versus the detected event number for both RD and DD. (**b**) Average translocation time ±SEM for DD and RD. (**c**) We calculated the average duration for groups of events at the beginning of our measurements (1-19) towards the end (58-79).

**Figure S10.**

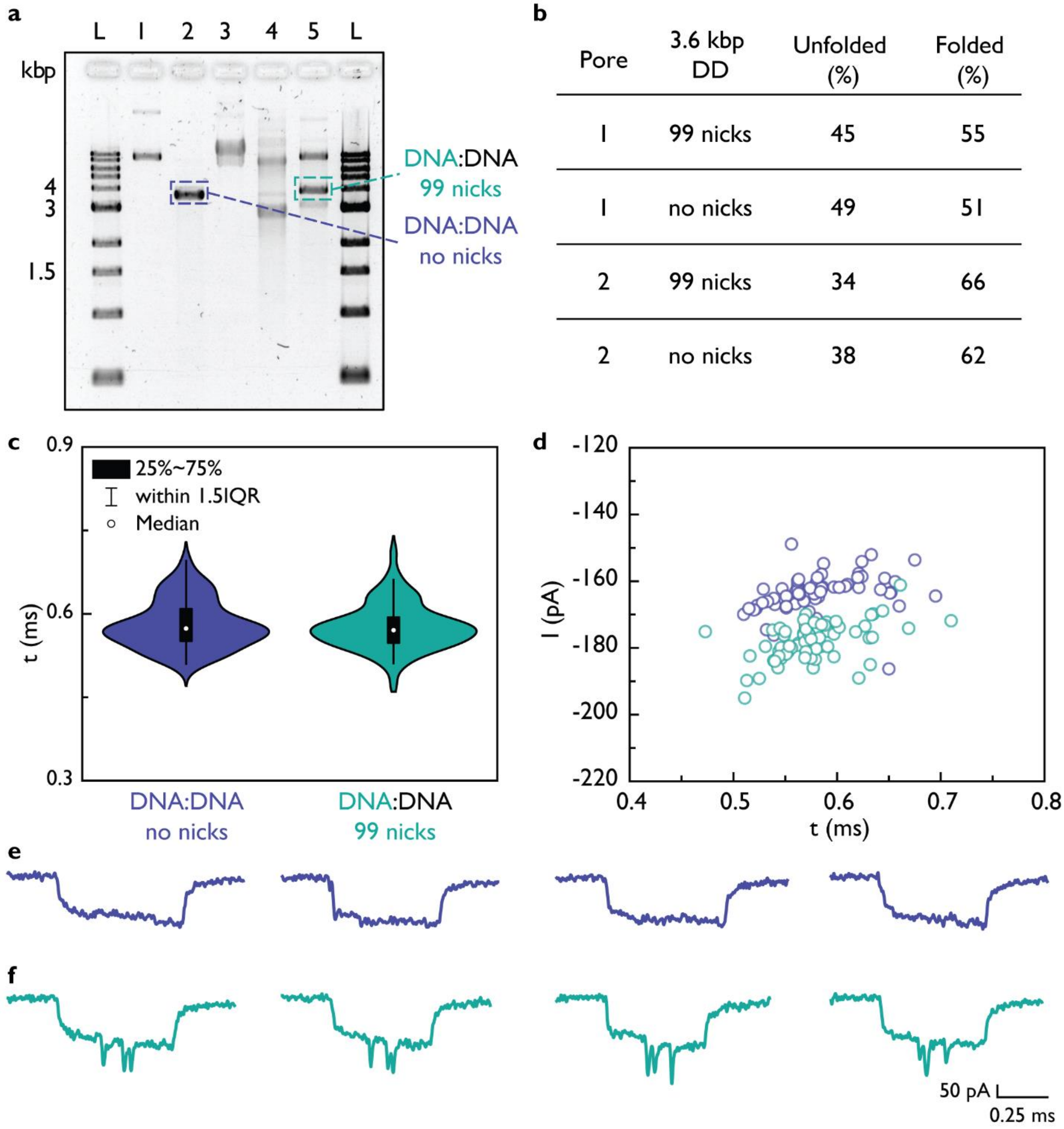


| Pore | 3.6 kbp DD | Unfolded (%) | Folded (%) |
|---|---|---|---|
| 1 | 99 nicks | 45 | 55 |
| 1 | no nicks | 49 | 51 |
| 2 | 99 nicks | 34 | 66 |
| 2 | no nicks | 38 | 62 |

**Figure S10.** Nanopore data for 3.6 kbp DD with 99 nicks and 3.6 kbp DD without nicks. (**a**) 1 % (w/v) agarose gel electrophoresis of preparation of linearized 3.6 kb ssDNA scaffold and 3.6 kbp dsDNA. L- 1 kbp ladder, NEB; 1 – 7.2kbp dsM13; 2 – 7.2 kbp dsM13 cut to yield 3.6 kbp DD without nicks; 3 – 7.2 kb ssM13; 4 - 7.2 kb ssM13 cut; 5 – 3.6 kbp DD with 99 nicks. (**b**) The percentage of unfolded (linear) and folded duplexes in our nanopore measurements. (**c**) The translocation time (τ) of 3.6 kbp DD no nicks (purple) and DD with 99 nicks (turquoise) were for both 0.58 ± 0.04 ms. (**d**) A scatter plot of the mean event current versus τ. The sample size was 100. Example nanopore events for DD without nicks and DD with 99 nicks are shown in (**e**) and (**f**), respectively.

**Figure S11.**

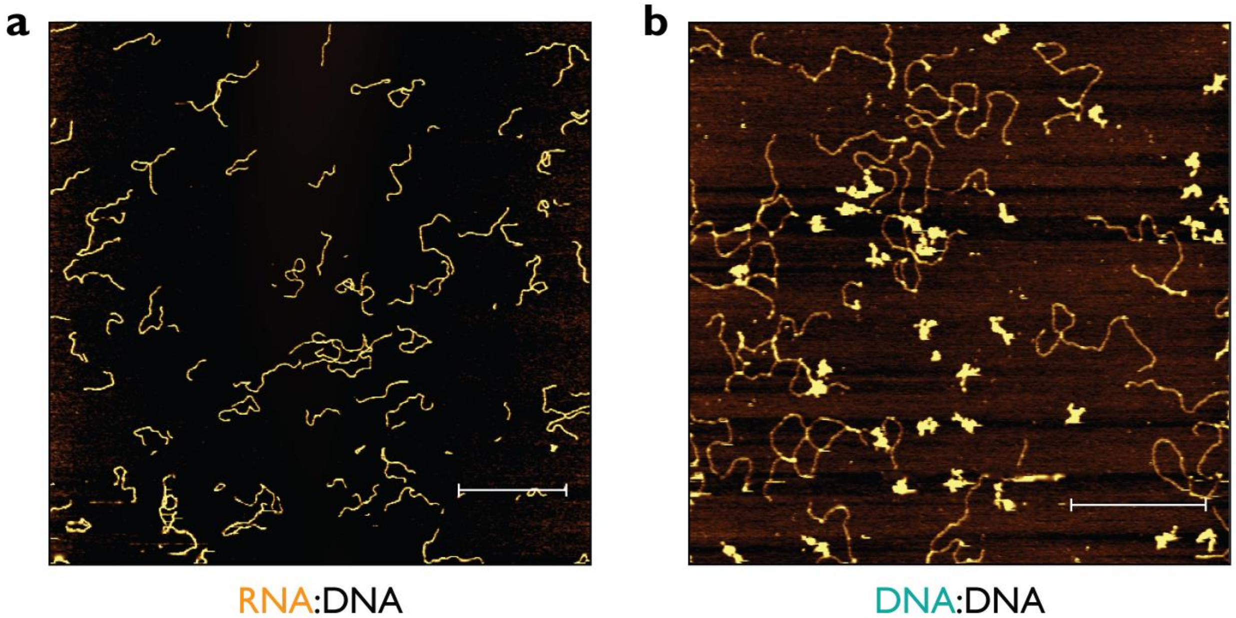


**Figure S11.** The 3.6 kbp RD and DD duplex length was verified using AFM imaging. (**a**) Example AFM image for RD duplexes. (**b**) Example AFM image for DD duplexes indicating the presence of coiled singe-cut ssM13 (brighter hank-like spots). The scale bars are 1 µm and 500 nm for (a) and (b), respectively.

**Figure S12.**

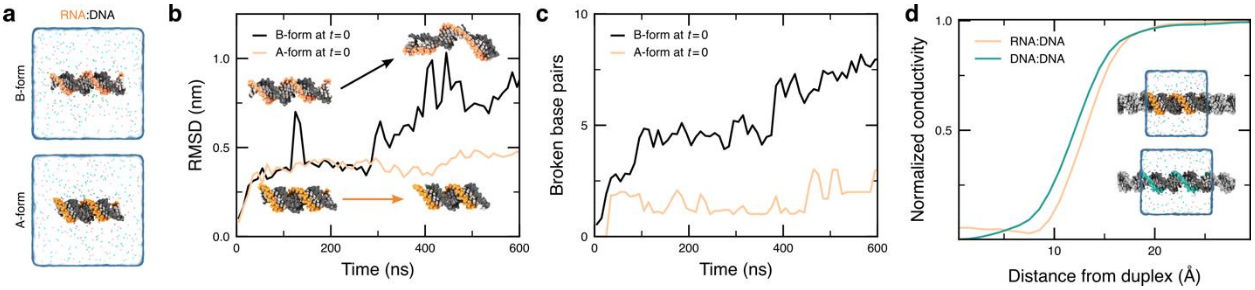


**Figure S12.** Simulated properties of RD duplexes. (**a**) All-atom MD simulation systems, each containing two helical turns of nucleic acid duplex solvated in 4 M LiCl electrolyte (cyan and magenta spheres depicted in the 9.4-Å thick slab centered on the duplex). The systems were constructed with idealized B- (top) and A-form (bottom) geometries. (**b**) Root mean square deviation of the phosphorus atoms from their initial idealized coordinates. (**c**) Number of broken base pairs during the simulations. (**d**) Dependence of ionic conductivity on the distance from RD and DD duplexes as observed during MD simulations of periodic helices.

**Figure S13.**

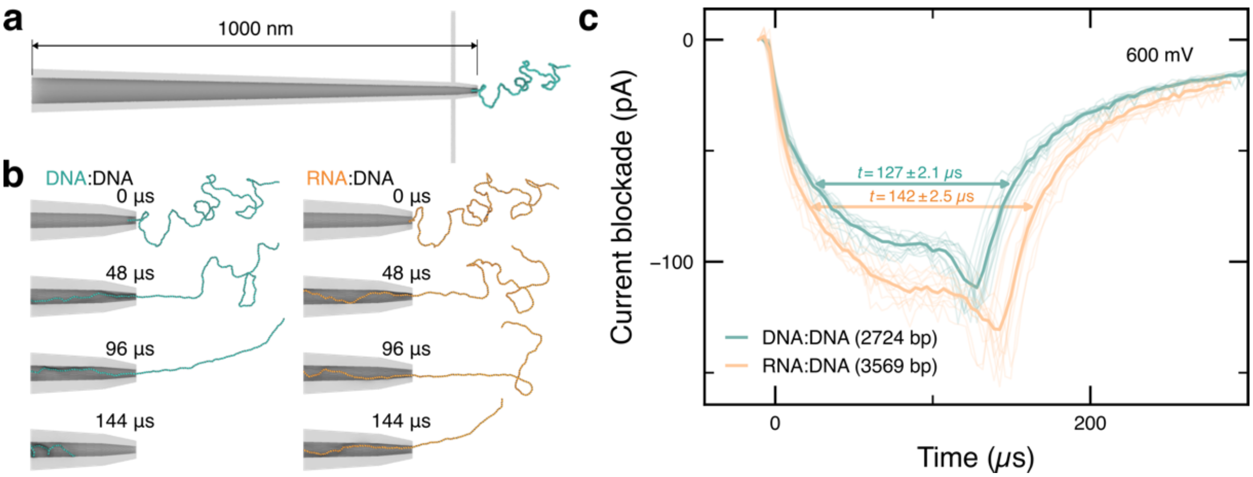


**Figure S13**. Coarse-grained Brownian dynamics simulations of same-contour-length DD and RD duplex translocation. (**a**) Initial configuration of a DD duplex threaded into a 1000 nm long conical nanopore having a 5-nm-radius inner aperture. (**b**) Snapshots depicting the translocation of DD and RD duplexes through the pore. The mobility and effective electric force acting on the duplexes were taken from the all-atom MD simulations. (**c**) Nanopore ionic current blockades computed from eight replicate DD and RD duplex translocation simulations. Each bold line depicts the current trace averaged over the eight replica simulations.

**Figure S14.**

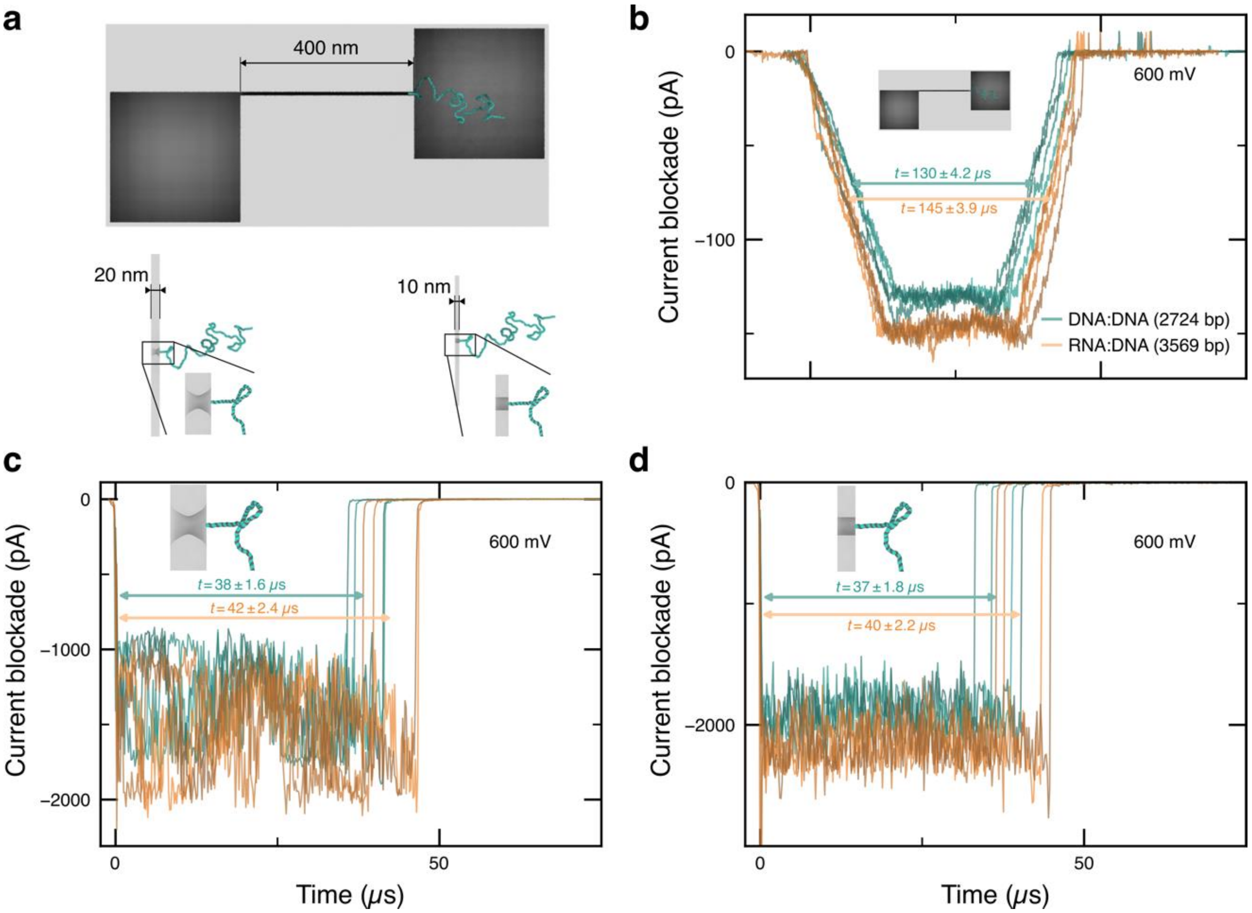


**Figure S14**. Coarse-grained Brownian dynamics simulations of same-contour-length DD and RD duplex translocation. (**a**) Initial configuration of DD duplexes in three model systems: a 400 nm long, 8 nm thick and 110 nm wide nanoslit (top); an hourglass-shaped nanopore of a 5-nm-radius constriction in a 20-nm-thick membrane (bottom left); and of a 5-nm-radius cylindrical nanopore in a 10-nm-thick membrane (bottom right). (**b-d**) Nanopore ionic current blockades computed for the four replicate DD and RD duplex translocation simulations, for each of the three systems illustrated in panel a.

**Figure S15.**

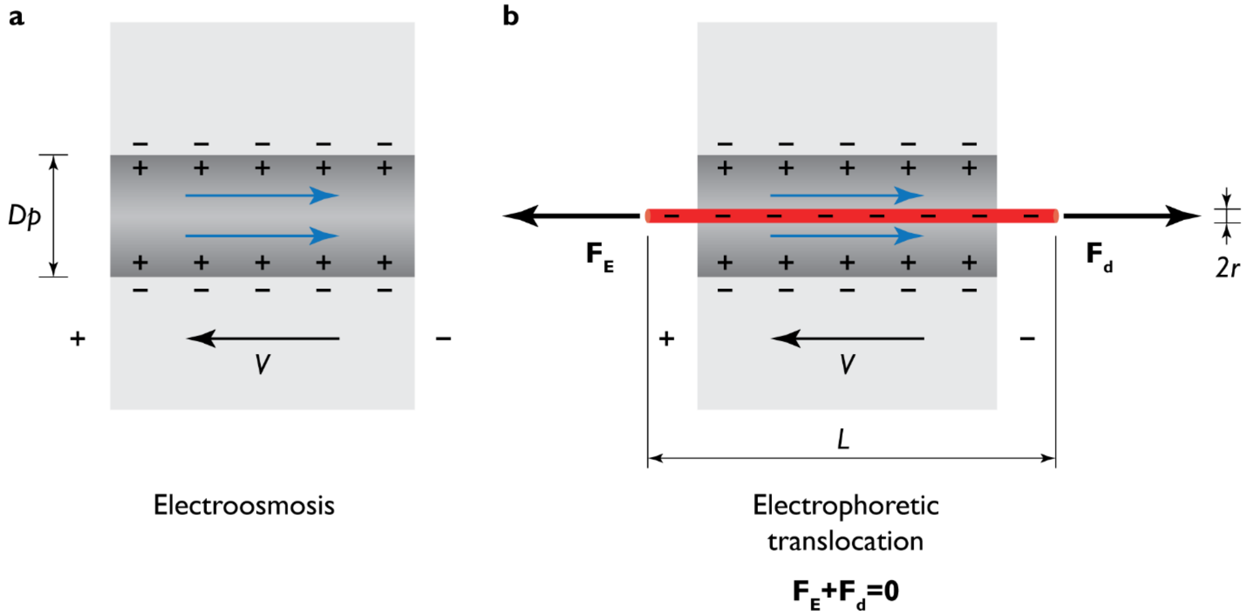


**Figure S15.** Translocation of nucleic acid duplex (red cylinder) through a nanopore (dark grey). (**a**) Voltage applied (*V*) across a negatively charged nanopore (with a diameter *Dp*) induces oppositely directed electroosmotic flows (blue arrows). (**b**) Nucleic acid duplex is represented by a negatively charged cylinder with length *L* and the radius *r*. The duplex is electrophoretically driven towards the positively charged electrode with the driving force $F_E$. The oppositely oriented force is the drag force $F_d$. Hence the total driving force $F_E$ on the duplex is always equal and opposite to the dissipative drag force $F_E + F_d = 0$ for a constant velocity *u*.

**Table S1.**

**Table S1.** Oligos used to assemble the 3.6 kbp MS2 RNA:DNA duplex '11100'. The overhang sequences to which the 3'- biotinylated imaging strand (5'-ACCACTAATGAGTGATATCC-3'TEG-biotin, HPLC purified, IDT) was bound are highlighted in red.

| Name | Sequence (5'→3') | Length (nt) |
|---|---|---|
| MS2_11100 | TGGGTGGTAACTAGCCAAGCAGCTAGTTACCAAATCGG | 38 |
| MS2_11100 | GAGAATCCCGGGTCCTCTCTTTAGGGGGAGGTCCCTGG | 38 |
| MS2_11100 | GCCGAAGCCCGCCCACCTTTCGGTGGAGCCGGACCGCT | 38 |
| MS2_11100 | TTCGCACCCGTGCTCTTTCGAGCACACCCACCCCGTTT | 38 |
| MS2_11100 | ACGGGGGTCCCTCGGTCAGCTACCGAGGAGAGCTCGCT | 38 |
| MS2_11100 | GGCCCACACTCCTGAGGGAATGTGGGAACCGGCGTTAG | 38 |
| MS2_11100 | CCACTCCGAAGTGCGTATAACGCGCACGCCGGCGGACT | 38 |
| MS2_11100 | TCATGCTGTCGGTGATTTCACCTCCAGTATGGAACCAC | 38 |
| MS2_11100 | GCTATGTAGCGACCACTGTCGTGCTTTTCGCTGAAGAA | 38 |
| MS2_11100 | CTTGCGTTCTCGAGCGATACGAGCAAGACGGAAACCCG | 38 |
| MS2_11100 | AGGTACGGGTATCCGCGAGCAGCCGCCCGTACGGAGTC | 38 |
| MS2_11100 | TTGGTGTATACCGAGACTGCCGTAGGCGGGCTGACTAC | 38 |
| MS2_11100 | GTAGTAGTCGGCAGCGAGGTCCGTCCCACCGAAGAACA | 38 |
| MS2_11100 | TCGAAGGCACCTGGGAGGAGAGCCGTACCCACACCTTA | 38 |
| MS2_11100 | TAGAGGCGTGGATCTGACATACCTCCGACAACTCCCCA | 38 |
| MS2_11100 | ACCCCGTAGCCGATTTAATATCAGCATCAGGGCGAAGA | 38 |
| MS2_11100 | GATTGTCAACAGGTTTCTTGATGTAAAACGGTTTGACA | 38 |
| MS2_11100 | TCGACACCACGGTAAAAGTGCGCGCCGCAGCTCTCGCG | 38 |
| MS2_11100 | AAAGAGCCCGGACACGAACGTTTTACGAAGATTCGGTT | 38 |
| MS2_11100 | TAAAACCGTAGTAGGCAAGTGCCTCTAGCACACGGGGT | 38 |
| MS2_11100 | GCAATCTCACTGGGACATATAATATCGTCCCCGTAGAT | 38 |
| MS2_11100 | GCCTATGGTTCCGGCGTTACCAAAATGGATTTGGGTCG | 38 |
| MS2_11100 | CTTTGACTATTGCCCAGAATATCATGGACTCTAGCTCA | 38 |
| MS2_11100 | AATGTGAACCCATTTCCCATTGTGGAAAATAGTTCCCA | 38 |
| MS2_11100 | TCGTATCGTCTCGCCATCTACGATTCCGTAGTGTGAGC | 38 |

| | | |
|---|---|---|
| MS2_11100 | GGATACGATCGAGATATGAATATAGCTCTGGTGGGAGA | 38 |
| MS2_11100 | AAACTCCACACCAGGCGATCGGAGATGGAATCGGATGC | 38 |
| MS2_11100 | AGACGATAAGTCTATCGTCGCAAGCGAACCATCTACGC | 38 |
| MS2_11100 | TGCCCTGCTGAGCCAGACGCTGGTTGATCGATTGATCA | 38 |
| MS2_11100 | TTCAGGTCTATACCAACGGATTTGAGCCGGCGTCTGAT | 38 |
| MS2_11100 | GAAAGCACCGACCCCTTTCTGGAGGTACATATTCATAT | 38 |
| MS2_11100 | CAGGCTCCTTACAGGCAGCCCGATCTATTTTATTATTC | 38 |
| MS2_11100 | TTCGGAACTGTAAACACTCCGTTCCCTACAACGAGCCT | 38 |
| MS2_11100 | AAATTCATATGACTCGTTATAGCGGACCGCGTGTCTGA | 38 |
| MS2_11100 | TCCACGGCGCACATTGGTCTCGGACCAATAGAGCCGCT | 38 |
| MS2_11100 | CTCAGAGCGCGGGGGGTAACGGTTGCTTGT | 30 |
| MS2_11100 | TCAGCGAACTTCTTGTAAGGCGCTGttt**ggatatcactcattagtggt** | 48 |
| MS2_11100 | CATCCTGCAACTTGTGCCCCATAGGAGCACCGTTGGAG | 38 |
| MS2_11100 | AACGTGCATTGCCCAAACAACGACGATCGGTAGCCAGA | 38 |
| MS2_11100 | GAGGAGGTTGCCAATAAGGCTACGGATGCTGGTTTGTA | 38 |
| MS2_11100 | AAACATCCGGATCCCATGACAAGGATTTGTCATGTAAG | 38 |
| MS2_11100 | AAACCTTCTCTATTTATCTGACC | 23 |
| MS2_11100 | GCGATCACCATTCGCCTCCCGTAGCttt**ggatatcactcattagtggt** | 48 |
| MS2_11100 | TTAGCGATAGCTAAGGTACGACGGGTCGCCTCGTCATT | 38 |
| MS2_11100 | ACCAGAACCTAAGGTCGGATGCTTTGTGAGCAATTCGT | 38 |
| MS2_11100 | CCCTTAAGTAAGCAATTGCTGTAAAGTCGTCACTGTGC | 38 |
| MS2_11100 | GGATCACCGCTTCCAGTAGCGACAGAAGCAATTGATTG | 38 |
| MS2_11100 | GTAAATTTCGAGAGAAAGATCGC | 23 |
| MS2_11100 | GAGGAAGATCAATACATAAAGAGTTttt**ggatatcactcattagtggt** | 48 |
| MS2_11100 | GAACTTCTTTGTTGTCTTCGACATGGGTAATCCTCATG | 38 |
| MS2_11100 | TTTGAATGGCCGGCGTCTATTAGTAGATGCCGGAGTTT | 38 |
| MS2_11100 | GCTGCGATTGCTGAGGGAATCGGGTTTCCATCTTTTAG | 38 |
| MS2_11100 | GAGACCTTGCATTGCCTTAACAATAAGCTCGCAGTCGG | 38 |
| MS2_11100 | AATTCGTAGCGAAAATTGGAATG | 23 |

| | | |
|---|---|---|
| MS2_11100 | GTTAGTTCCATATTTAAGTACGAAC | 25 |
| MS2_11100 | GCCATGCGGCTACAGGAAGCTCTACACCACCAACAGTC | 38 |
| MS2_11100 | TGGGTTGCCACTTTAGGCACCTCGACTTTGATGGTGTA | 38 |
| MS2_11100 | TTTGCGATTCTGCGCAGAGCTCTGACGAACGCTACAGG | 38 |
| MS2_11100 | TTACTTTGTAAGCCTGTGAACGCGAGTTAGAGCTGATC | 38 |
| MS2_11100 | CATTCAGCGACCCCGTTAGCGAA | 23 |
| MS2_11100 | GTTGCTTGGGGCGACAGTCACGTCG | 25 |
| MS2_11100 | CCAGTTCCGCCATTGTCGACGAGAACGAACTGAGTAAA | 38 |
| MS2_11100 | GTTAGAAGCCATGCTTCAAACTCCGGTTGAGGGCTCTA | 38 |
| MS2_11100 | TCTAGAGAGCCGTTGCCTGATTAATGCTAACGCATCTA | 38 |
| MS2_11100 | AGGTATGGACCATCGAGAAAGGAGACTTTACGTACGCG | 38 |
| MS2_11100 | CCAGTTGTTGGCCATACGGATTGTACCCCTCGATGCAT | 38 |
| MS2_11100 | GGCTGAGATTTGGGCCTTAGCAGTGCCCTGTCTCTCCA | 38 |
| MS2_11100 | CAGTCCACCCGTAGGGAGCGTCAACGCTTATGATGGAC | 38 |
| MS2_11100 | TCACCCGTTATTACGTCAGTAACTGTTCCTGACATGTA | 38 |
| MS2_11100 | GGAGCATCCCACGGGGGCCGTAAGGCCCTCGAGCATGT | 38 |
| MS2_11100 | TACCTACAGGTAGGAGCCAGTCGACAACGAATGAGAAA | 38 |
| MS2_11100 | GGCACCTTTTCCCACACTATACCTAGTGGGTTCAAGAT | 38 |
| MS2_11100 | ACCTAGAGACGACAACCATGCCAAACGTGCATCGTTTA | 38 |
| MS2_11100 | TGTAAAACCATATCACGATACGTCGCGATATGTTGCAC | 38 |
| MS2_11100 | GTTGTCTGGAAGTTTGCAGCTGGATACGACAGACGGCC | 38 |
| MS2_11100 | ATCTAACTTGATGTTAGTACCGACCTGACGTACGGCTC | 38 |
| MS2_11100 | TCATAGGAAGAAACTCTTGAAGGTGAACCTTCGTAAGC | 38 |
| MS2_11100 | ATCTCATATGCACCCTGGATATCACTCATTAGTGGTAA | 38 |
| MS2_11100 | CCAACCGAACTGCAACTCCAACCACCTGCCGGCCACGT | 38 |
| MS2_11100 | GTTTTGATCGAAACTTTCGATCTTCGTTTAGGGCAAGG | 38 |
| MS2_11100 | TAGCGGAGCGCCTGGCGCCAATTACCGCGACGAGCGGC | 38 |
| MS2_11100 | AGTGTACGCCTTCACGAGCGCAATGGTTTGCGTCGCGA | 38 |
| MS2_11100 | GTTGTGAGGCTGTCGACCTGGCCTCTGCTAAAGCAACA | 38 |

| | | |
|---|---|---|
| MS2_11100 | CCAAGGTTAAAATTACCCTGGGTGACCTTTTGCAGGAC | 38 |
| MS2_11100 | TTCGGTCGACGCCCGGTTCGCAACGTTCTGCGGCACTT | 38 |
| MS2_11100 | CGATGTAAGTCAAGTTTTGGCTTACAGGGAAGAGGCTG | 38 |
| MS2_11100 | TAGCAGGAGCGTGCGTCGAGGGAGAAGCCGAAACCGGC | 38 |
| MS2_11100 | TTTCTCCTCGTACGGGCGACCCCACGATGACCCACTTC | 38 |
| MS2_11100 | GCTTGTAGGCACCTTGATCTATCGATGTGACACTTAAC | 38 |
| MS2_11100 | GCCCCCCGTGAATACGGAGAGGGGTAGTGCCACTGTTT | 38 |
| MS2_11100 | CGTTTTGGCCCCAGTCGAGTTAAAACGACCGGGAGTCC | 38 |
| MS2_11100 | AGTTCGAACGATATTTTAAAGAGAATGAGTTATCTTCA | 38 |
| MS2_11100 | GTCTCACCGTCCGCGTAAACGCGAACGGAGGGGACGAA | 38 |
| MS2_11100 | GGTCTCGTTCTCCCTATCAAGGGTACTAAAAGCTCGCA | 38 |
| MS2_11100 | CAGGTCAAACCTCCTAGGAATGGAATTCCGGCTACCTA | 38 |
| MS2_11100 | CAGCGATAGCCATGGTAGCGTCTCGCTAAAGACATTAA | 38 |
| MS2_11100 | AAATGGCATTAGCTCGACAGGAAGTTG | 27 |
| MS2_11100 | AGCAGGACCCCGAAAGGGGTCCCACCC | 27 |

**Table S2.**

**Table S2.** Oligos used to assemble the 3.6 kbp DNA:DNA duplex '11001'. The overhang sequences to which the 3'- biotinylated imaging strand (5'-ACCACTAATGAGTGATATCC-3'TEG-biotin, HPLC purified, IDT) was bound are highlighted in red.

| Name | Sequence (5’→3’) | Length (nt) |
|---|---|---|
| M13_11001 | TGCCCGTATAAACAGTTAATGCCCCCTGCCTATTTCGG | 38 |
| M13_11001 | AACCTATTATTCTGAAACATGAAAGTATTAAGAGGCTG | 38 |
| M13_11001 | AGACTCCTCAAGAGAAGGATTAGGATTAGCGGGGTTTT | 38 |
| M13_11001 | GCTCAGTACCAGGCGGATAAGTGCCGTCGAGAGGGTTG | 38 |
| M13_11001 | ATATAAGTATAGCCCGGAATAGGTGTATCACCGTACTC | 38 |
| M13_11001 | AGGAGGTTTAGTACCGCCACCCTCAGAACCGCCACCCT | 38 |
| M13_11001 | CAGAACCGCCACCCTCAGAGCCACCACCCTCATTTTCA | 38 |
| M13_11001 | GGGATAGCAAGCCCAATAGGAACCCATGTACCGTAACA | 38 |
| M13_11001 | CTGAGTTTCGTCACCAGTACAAACTACAACGCCTGTAG | 38 |
| M13_11001 | CATTCCACAGACAGCCCTCATAGTTAGCGTAACGATCT | 38 |
| M13_11001 | AAAGTTTTGTCGTCTTTCCAGACGTTAGTAAATGAATT | 38 |
| M13_11001 | TTCTGTATGGGATTTTGCTAAACAACTTTCAACAGTTT | 38 |
| M13_11001 | CAGCGGAGTGAGAATAGAAAGGAACAACTAAAGGAATT | 38 |
| M13_11001 | GCGAATAATAATTTTTTCACGTTGAAAATCTCCAAAAA | 38 |
| M13_11001 | AAAGGCTCCAAAAGGAGCCTTTAATTGTATCGGTTTAT | 38 |
| M13_11001 | CAGCTTGCTTTCGAGGTGAATTTCTTAAACAGCTTGAT | 38 |
| M13_11001 | ACCGATAGTTGCGCCGACAATGACAACAACCATCGCCC | 38 |
| M13_11001 | ACGCATAACCGATATATTCGGTCGCTGAGGCTTGCAGG | 38 |
| M13_11001 | GAGTTAAAGGCCGCTTTTGCGGGATCGTCACCCTCAGC | 38 |
| M13_11001 | AGCGAAAGACAGCATCGGAACGAGGGTAGCAACGGCTA | 38 |
| M13_11001 | CAGAGGCTTTGAGGACTAAAGACTTTTTCATGAGGAAG | 38 |
| M13_11001 | TTTCCATTAAACGGGTAAAATACGTAATGCCACTACGA | 38 |
| M13_11001 | AGGCACCAACCTAAAACGAAAGAGGCAAAAGAATACAC | 38 |
| M13_11001 | TAAAACACTCATCTTTGACCCCCAGCGATTATACCAAG | 38 |
| M13_11001 | CGCGAAACAAAGTACAACGGAGATTTGTATCATCGCCT | 38 |

| | | |
|---|---|---|
| M13_11001 | GATAAATTGTGTCGAAATCCGCGACCTGCTCCATGTTA | 38 |
| M13_11001 | CTTAGCCGGAACGAGGCGCAGACGGTCAATCATAAGGG | 38 |
| M13_11001 | AACCGAACTGACCAACTTTGAAAGAGGACAGATGAACG | 38 |
| M13_11001 | GTGTACAGACCAGGCGCATAGGCTGGCTGACCTTCATC | 38 |
| M13_11001 | AAGAGTAATCTTGACAAGAACCGGATATTCATTACCCA | 38 |
| M13_11001 | AATCAACGTAACAAAGCTGCTCATTCAGTGAATAAGGC | 38 |
| M13_11001 | TTGCCCTGACGAGAAACACCAGAACGAGTAGTAAATTG | 38 |
| M13_11001 | GGCTTGAGATGGTTTAATTTCAACTTTAATCATTGTGA | 38 |
| M13_11001 | ATTACCTTATGCGATTTTAAGAACTGGCTCATTATACC | 38 |
| M13_11001 | AGTCAGGACGTTGGGAAGAAAAATCTACGTTAATAAAA | 38 |
| M13_11001 | CGAACTAACGGAACAACATTATTACAG | 27 |
| M13_11001 | GTAGAAAGATTCATCAGTTGAGATTTAG | 28 |
| M13_11001 | GAATACCACATTCAACTAATGCAGAttt**ggatatcactcattagtggt** | 48 |
| M13_11001 | TACATAACGCCAAAAGGAATTACGAGGCATAGTAAGAG | 38 |
| M13_11001 | CAACACTATCATAACCCTCGTTTACCAGACGACGATAA | 38 |
| M13_11001 | AAACCAAAATAGCGAGAGGCTTTTGCAAAAGAAGTTTT | 38 |
| M13_11001 | GCCAGAGGGGGTAATAGTAAAATGTTTAGACTGGATAG | 38 |
| M13_11001 | CGTCCAATACTGCGGAATCGTCA | 23 |
| M13_11001 | TAAATATTCATTGAATCCCCCTCAAttt**ggatatcactcattagtggt** | 48 |
| M13_11001 | ATGCTTTAAACAGTTCAGAAAACGAGAATGACCATAAA | 38 |
| M13_11001 | TCAAAAATCAGGTCTTTACCCTGACTATTATAGTCAGA | 38 |
| M13_11001 | AGCAAAGCGGATTGCATCAAAAAGATTAAGAGGAAGCC | 38 |
| M13_11001 | CGAAAGACTTCAAATATCGCGTTTTAATTCGAGCTTCA | 38 |
| M13_11001 | AAGCGAACCAGACCGGAAGCAAA | 23 |
| M13_11001 | CTCCAACAGGTCAGGATTAGAGAGT | 25 |
| M13_11001 | ACCTTTAATTGCTCCTTTTGATAAGAGGTCATTTTTGC | 38 |
| M13_11001 | GGATGGCTTAGAGCTTAATTGCTGAATATAATGCTGTA | 38 |
| M13_11001 | GCTCAACATGTTTTAAATATGCAACTAAAGTACGGTGT | 38 |
| M13_11001 | CTGGAAGTTTCATTCCATATAACAGTTGATTCCCAATT | 38 |

| | | |
|---|---|---|
| M13_11001 | CTGCGAACGAGTAGATTTAGTTT | 23 |
| M13_11001 | GACCATTAGATACATTTCGCAAATG | 25 |
| M13_11001 | GTCAATAACCTGTTTAGCTATATTTTCATTTGGGGCGC | 38 |
| M13_11001 | GAGCTGAAAAGGTGGCATCAATTCTACTAATAGTAGTA | 38 |
| M13_11001 | GCATTAACATCCAATAAATCATACAGGCAAGGCAAAGA | 38 |
| M13_11001 | ATTAGCAAAATTAAGCAATAAAGCCTCAGAGCATAAAG | 38 |
| M13_11001 | CTAAATCGGTTGTACCAAAAACA | 23 |
| M13_11001 | TTATGACCCTGTAATACTTTTGCGGttt**ggatatcactcattagtggt** | 48 |
| M13_11001 | GAGAAGCCTTTATTTCAACGCAAGGATAAAAATTTTTA | 38 |
| M13_11001 | GAACCCTCATATATTTTAAATGCAATGCCTGAGTAATG | 38 |
| M13_11001 | TGTAGGTAAAGATTCAAAAGGGTGAGAAAGGCCGGAGA | 38 |
| M13_11001 | CAGTCAAATCACCATCAATATGATATTCAACCGTTCTA | 38 |
| M13_11001 | GCTGATAAATTAATGCCGGAGAGGGTAGCTATTTTTGA | 38 |
| M13_11001 | GAGATCTACAAAGGCTATCAGGTCATTGCCTGAGAGTC | 38 |
| M13_11001 | TGGAGCAAACAAGAGAATCGATGAACGGTAATCGTAAA | 38 |
| M13_11001 | ACTAGCATGTCAATCATATGTACCCCGGTTGATAATCA | 38 |
| M13_11001 | GAAAAGCCCCAAAAACAGGAAGATTGTATAAGCAAATA | 38 |
| M13_11001 | TTTAAATTGTAAACGTTAATATTTTGTTAAAATTCGCA | 38 |
| M13_11001 | TTAAATTTTTGTTAAATCAGCTCATTTTTTAACCAATA | 38 |
| M13_11001 | GGAACGCCATCAAAAATAATTCGCGTCTGGCCTTCCTG | 38 |
| M13_11001 | TAGCCAGCTTTCATCAACATTAAATGTGAGCGAGTAAC | 38 |
| M13_11001 | AACCCGTCGGATTCTCCGTGGGAACAAACGGCGGATTG | 38 |
| M13_11001 | ACCGTAATGGGATAGGTCACGTTGGTGTAGATGGGCGC | 38 |
| M13_11001 | ATCGTAACCGTGCATCTGCCAGTTTGAGGGGACGACGA | 38 |
| M13_11001 | CAGTATCGGCCTCAGGAAGATCGCACTCCAGCCAGCTT | 38 |
| M13_11001 | TCCGGCACCGCTTCTGGTGCCGGAAACCAGGCAAAGCG | 38 |
| M13_11001 | CCATTCGCCATTCAGGCTGCGCAACTGTTGGGAAGGGC | 38 |
| M13_11001 | GATCGGTGCGGGCCTCTTCGCTATTACGCCAGCTGGCG | 38 |
| M13_11001 | AAAGGGGGATGTGCTGCAAGGCGATTAAGTTGGGTAAC | 38 |

| | | |
|---|---|---|
| M13_11001 | GCCAGGGTTTTCCCAGTCACGACGTTGTAAAACGACGG | 38 |
| M13_11001 | CCAGTGCCAAGCTTGCATGCCTGCAGGTCGACTCTAGA | 38 |
| M13_11001 | GGATCCCCGGGTACCGAGCTCGAATTCGTAATCATGGT | 38 |
| M13_11001 | CATAGCTGTTTCCTGTGTGAAATTGTTATCCGCTCACA | 38 |
| M13_11001 | ATTCCACACAACATACGAGCCGGAAGCATAAAGTGTAA | 38 |
| M13_11001 | AGCCTGGGGTGCCTAATGAGTGAGCTAACTCACATTAA | 38 |
| M13_11001 | TTGCGTTGCGCTCACTGCCCGCTTTCCAGTCGGGAAAC | 38 |
| M13_11001 | CTGTCGTGCCAGCTGCATTAATGAATCGGCCAACGCGC | 38 |
| M13_11001 | GGGGAGAGGCGGTTTGCGTATTGGGCGCCAGGGTGGTT | 38 |
| M13_11001 | TTTCTTTTCACCAGTGAGACGGGCAACAGCTGATTGCC | 38 |
| M13_11001 | CTTCACCGCCTGGCCCTGAGAGAGTTGCAGCAAGCGGT | 38 |
| M13_11001 | CCACGCTGGTTTGCCCCAGCAGGCGAAAATCCTGTTTG | 38 |
| M13_11001 | ATGGTGGTTCCGAAATCGGCAAAATCCCTTATAAATCA | 38 |
| M13_11001 | AAAGAATAGCCCGAGATAGGGTTGAGTGTTGTTCCAGT | 38 |
| M13_11001 | TTGGAACAAGAGTCCACTATTAAAGAACGTGGACTCCA | 38 |
| M13_11001 | ACGTCAAAGGGCGAAAAACCG | 21 |
| M13_11001 | TCTATCAGGGCGATGGCCCAC | 21 |

**Table S3.**

**Table S3.** Parameters for nanopores used for measuring of 3.6 kbp RNA:DNA and 3.6 kbp DNA:DNA samples.

| Pore number | Sample | Current and RMS noise at 600 mV | Current / voltage curve (IV curve) |
|---|---|---|---|
| **1** | **3.6 kbp RNA:DNA, 3.6 kbp DNA:DNA** | **8.3 nA**<br>**6 pA** | 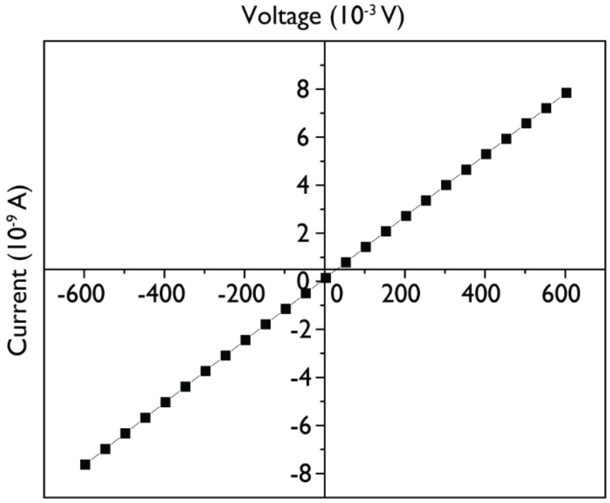  |
| **2** | | **6.5 nA**<br>**6 pA** | 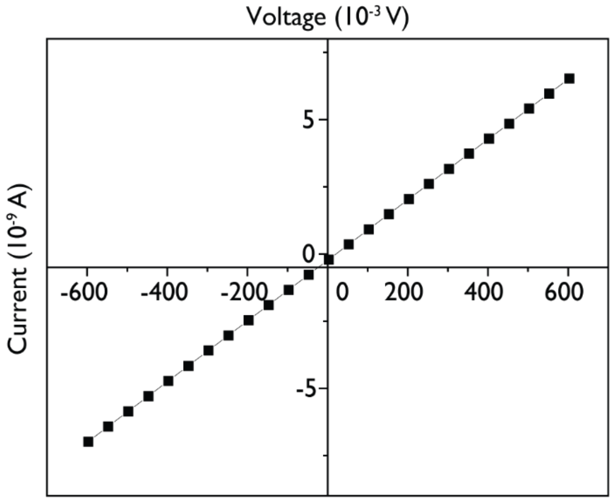  |

| | | |
|---|---|---|
| **3** | **7.9 nA**<br>**6 pA** | 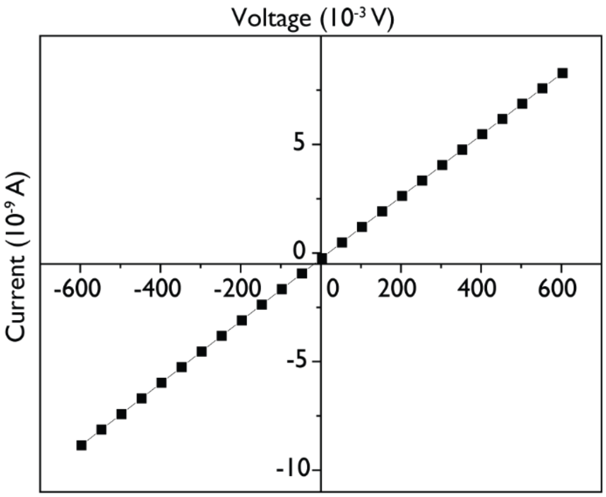 |

**Table S4.**

**Table S4.** Oligos used to assemble the 2.7 kbp DNA:DNA duplex '11001'. The overhang sequences to which the 3'- biotinylated imaging strand (5'-ACCACTAATGAGTGATATCC-3'TEG-biotin, HPLC purified, IDT) was bound are highlighted in red.

| Name | Sequence (5'→3') | Length (nt) |
|---|---|---|
| M13_2.7 kb_11001 | CAACGTCAAAGGGCGAAAAACCGTCTATCAGGGCGATGGC | 40 |
| M13_2.7 kb_11001 | CCACTACGTGAACCATCACCCAAATCAAGTTTTTTGGGGT | 40 |
| M13_2.7 kb_11001 | CGAGGTGCCGTAAAGCACTAAATCGGAACCCTAAAGGGAG | 40 |
| M13_2.7 kb_11001 | CCCCCGATTTAGAGCTTGACGGGGAAAGCCGGCGAACGTG | 40 |
| M13_2.7 kb_11001 | GCGAGAAAGGAAGGGAAGAAAGCGAAAGGAGCGGGCGCTA | 40 |
| M13_2.7 kb_11001 | GGGCGCTGGCAAGTGTAGCGGTCACGCTGCGCGTAACCAC | 40 |
| M13_2.7 kb_11001 | CACACCCGCCGCGCTTAATGCGCCGCTACAGGGCGCGTAC | 40 |
| M13_2.7 kb_11001 | TATGGTTGCTTTGACGAGCACGTATAACGTGCTTTCCTCG | 40 |
| M13_2.7 kb_11001 | TTAGAATCAGAGCGGGAGCTAAACAGGAGGCCGATTAAAG | 40 |
| M13_2.7 kb_11001 | GGATTTTAGACAGGAACGGTACGCCAGAATCCTGAGAAGT | 40 |
| M13_2.7 kb_11001 | GTTTTTATAATCAGTGAGGCCACCGAGTAAAAGAGTCTGT | 40 |
| M13_2.7 kb_11001 | CCATCACGCAAATTAACCGTTGTAGCAATACTTCTTTGAT | 40 |
| M13_2.7 kb_11001 | TAGTAATAACATCACTTGCCTGAGTAGAAGAACTCAAACT | 40 |
| M13_2.7 kb_11001 | ATCGGCCTTGCTGGTAATATCCAGAACAATATTACCGCCA | 40 |
| M13_2.7 kb_11001 | GCCATTGCAACAGGAAAAACGCTCATGGAAATACCTACAT | 40 |
| M13_2.7 kb_11001 | TTTGACGCTCAATCGTCTGAAATGGATTATTTACATTGGC | 40 |
| M13_2.7 kb_11001 | AGATTCACCAGTCACACGACCAGTAATAAAAGGGACATTC | 40 |
| M13_2.7 kb_11001 | TGGCCAACAGAGATAGAACCCTTCTGACCTGAAAGCGTAA | 40 |
| M13_2.7 kb_11001 | GAATACGTGGCACAGACAATATTTTTGAATGGCTATTAGT | 40 |
| M13_2.7 kb_11001 | CTTTAATGCGCGAACTGATAGCCCTAAAACATCGCCATTA | 40 |
| M13_2.7 kb_11001 | AAAATACCGAACGAACCACCAGCAGAAGATAAAACAGAGG | 40 |
| M13_2.7 kb_11001 | TGAGGCGGTCAGTATTAACACCGCCTGCAACAGTGCCACG | 40 |
| M13_2.7 kb_11001 | CTGAGAGCCAGCAGCAAATGAAAAATCT | 28 |
| M13_2.7 kb_11001 | AAAGCATCACCTTGCTGAACCTCAAATAT | 29 |

| | | |
|---|---|---|
| M13_2.7 kb_11001 | CAAACCCTCAATCAATATCTGGTCAttt**ggatatcactcattagtggt** | 48 |
| M13_2.7 kb_11001 | GTTGGCAAATCAACAGTTGAAAGGAATTGAGGAAGGTTAT | 40 |
| M13_2.7 kb_11001 | CTAAAATATCTTTAGGAGCACTAACAACTAATAGATTAGA | 40 |
| M13_2.7 kb_11001 | GCCGTCAATAGATAATACATTTGAGGATTTAGAAGTATTA | 40 |
| M13_2.7 kb_11001 | GACTTTACAAACAATTCGACAACTCGT | 27 |
| M13_2.7 kb_11001 | ATTAAATCCTTTGCCCGAACGTTATTAA | 28 |
| M13_2.7 kb_11001 | TTTTAAAAGTTTGAGTAACATTATCttt**ggatatcactcattagtggt** | 48 |
| M13_2.7 kb_11001 | ATTTTGCGGAACAAAGAAACCACCAGAAGGAGCGGAATTA | 40 |
| M13_2.7 kb_11001 | TCATCATATTCCTGATTATCAGATGATGGCAATTCATCAA | 40 |
| M13_2.7 kb_11001 | TATAATCCTGATTGTTTGGATTATACTTCTGAATAATGGA | 40 |
| M13_2.7 kb_11001 | AGGGTTAGAACCTACCATATCAAAATTATTTGCACGTAAA | 40 |
| M13_2.7 kb_11001 | ACAGAAATAAAGAAATTGCGTAGATTTTCAGGTTTAACGT | 40 |
| M13_2.7 kb_11001 | CAGATGAATATACAGTAACAGTACCTTTTACATCGGGAGA | 40 |
| M13_2.7 kb_11001 | AACAATAACGGATTCGCCTGATTGCTTTGAATACCAAGTT | 40 |
| M13_2.7 kb_11001 | ACAAAATCGCGCAGAGGCGAATTATTCATTTCAATTACCT | 40 |
| M13_2.7 kb_11001 | GAGCAAAAGAAGATGATGAAACAAACATCAAGAAAACAAA | 40 |
| M13_2.7 kb_11001 | ATTAATTACATTTAACAATTTCATTTGAATTACCTTTTTT | 40 |
| M13_2.7 kb_11001 | AATGGAAACAGTACATAAATCAATATATGTGAGTGAATAA | 40 |
| M13_2.7 kb_11001 | CCTTGCTTCTGTAAATCGTCGCTATTAATTAATTTTCCCT | 40 |
| M13_2.7 kb_11001 | TAGAATCCTTGAAAACATAGCGATAGCTTAGATTAAGACG | 40 |
| M13_2.7 kb_11001 | CTGAGAAGAGTCAATAGTGAATTTATC | 27 |
| M13_2.7 kb_11001 | AAAATCATAGGTCTGAGAGACTACCTTT | 28 |
| M13_2.7 kb_11001 | TTAACCTCCGGCTTAGGTTGGGTTAttt**ggatatcactcattagtggt** | 48 |
| M13_2.7 kb_11001 | TATAACTATATGTAAATGCTGATGCAAATCCAATCGCAAG | 40 |
| M13_2.7 kb_11001 | ACAAAGAACGCGAGAAAACTTTTTCAAATATATTTTAGTT | 40 |
| M13_2.7 kb_11001 | AATTTCATCTTCTGACCTAAATTTAATGGTTTGAAATACC | 40 |
| M13_2.7 kb_11001 | GACCGTGTGATAAATAAGGCGTTAAATAAGAATAAACACC | 40 |
| M13_2.7 kb_11001 | GGAATCATAATTACTAGAAAAAGCCTGTTTAGTATCATAT | 40 |
| M13_2.7 kb_11001 | GCGTTATACAAATTCTTACCAGTATAAAGCCAACGCTCAA | 40 |

| | | |
|---|---|---|
| M13_2.7 kb_11001 | CAGTAGGGCTTAATTGAGAATCGCCATATTTAACAACGCC | 40 |
| M13_2.7 kb_11001 | AACATGTAATTTAGGCAGAGGCATTTTCGAGCCAGTAATA | 40 |
| M13_2.7 kb_11001 | AGAGAATATAAAGTACCGACAAAAGGTAAAGTAATTCTGT | 40 |
| M13_2.7 kb_11001 | CCAGACGACGACAATAAACAACATGTTCAGCTAATGCAGA | 40 |
| M13_2.7 kb_11001 | ACGCGCCTGTTTATCAACAATAGATAAGTCCTGAACAAGA | 40 |
| M13_2.7 kb_11001 | AAAATAATATCCCATCCTAATTTACGAGCATGTAGAAACC | 40 |
| M13_2.7 kb_11001 | AATCAATAATCGGCTGTCTTTCCTTATCATTCCAAGAACG | 40 |
| M13_2.7 kb_11001 | GGTATTAAACCAAGTACCGCACTCATCGAGAACAAGCAAG | 40 |
| M13_2.7 kb_11001 | CCGTTTTTATTTTCATCGTAGGAATCATTACCGCGCCCAA | 40 |
| M13_2.7 kb_11001 | TAGCAAGCAAATCAGATATAGAAGGCTTATCCGGTATTCT | 40 |
| M13_2.7 kb_11001 | AAGAACGCGAGGCGTTTTAGCGAACCTCCCGACTTGCGGG | 40 |
| M13_2.7 kb_11001 | AGGTTTTGAAGCCTTAAATCAAGATTAGTTGCTATTTTGC | 40 |
| M13_2.7 kb_11001 | ACCCAGCTACAATTTTATCCTGAATCTTACCAACGCTAAC | 40 |
| M13_2.7 kb_11001 | GAGCGTCTTTCCAGAGCCTAATTTGCCAGTTACAAAATAA | 40 |
| M13_2.7 kb_11001 | ACAGCCATATTATTTATCCCAATCCAAATAAGAAACGATT | 40 |
| M13_2.7 kb_11001 | TTTTGTTTAACGTCAAAAATGAAAATAGCAGCCTTTACAG | 40 |
| M13_2.7 kb_11001 | AGAGAATAACATAAAAACAGGGAAGCGCATTAGACGGGAG | 40 |
| M13_2.7 kb_11001 | AATTAACTGAACACCCTGAACAAAGTCAGAGGGTAATTGAGC | 42 |

**Table S5.**

**Table S5.** Parameters for nanopores used for measuring of 3.6 kbp RNA:DNA and 2.7 kbp DNA:DNA samples.

| Pore number | Sample | Current and RMS noise at 600 mV | Current / voltage curve (IV curve) |
|---|---|---|---|
| 1 | 3.6 kbp RNA:DNA, 2.7 kbp DNA:DNA | 9.3 nA, 6 pA | 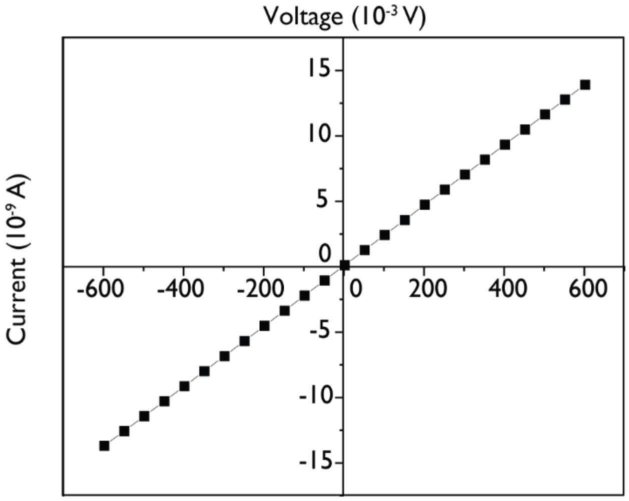  |
| 2 | | 13.9 nA, 7 pA | 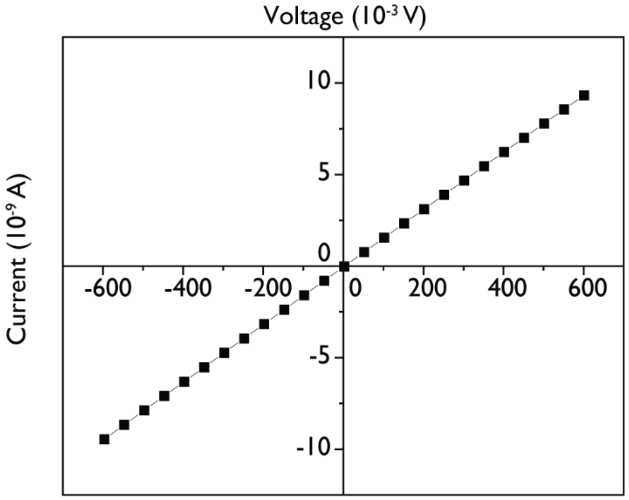  |

| | | |
|---|---|---|
| **3** | **8.1 nA,<br>7 pA** | 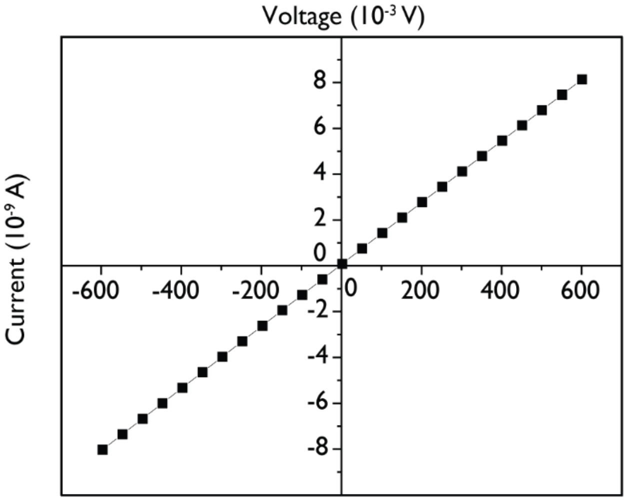 |
| **4** | **7.8 nA,<br>7 pA** | 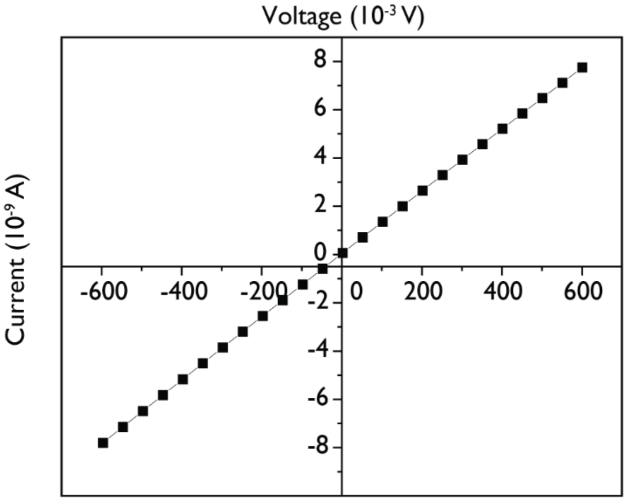 |

**Table S6.**

**Table S6.** Force ratios for 3.6 kbp RD and DD using nanopore measurements.

| **Pore #** | **Current at 600 mV** | **Data shown in** | **Event number** | **Force ratio ($F_{RD}/F_{DD}$)** |
|---|---|---|---|---|
| 1 | 8.3 nA | Fig.1 | 80 | 1.12 |
| 2 | 6.5 nA | Fig.S5a,b | 589 | 0.97 |
| 3 | 7.9 nA | Fig.S5c,d | 118 | 1.02 |
| 4 | 7.2 nA | Fig.S5e,f | 82 | 0.94 |
| 5 | 7.8 nA | Fig.S5g,h | 100 | 0.92 |
| 6 | 9.6 nA | Fig.S5i,j | 77 | 0.95 |
| 7 | 10.7 nA | Fig.S5k,l | 97 | 0.93 |
| Force ratio average (for pores 1-7) | | 0.98 | | |
| Force ratio standard error ± (for pores 1-7) | | 0.07 | | |

**Table S7.**

**Table S7.** Parameters for nanopores used for measuring of 3.6 kbp DNA:DNA with 99 nicks and without nicks (Figure S10).

| Pore number | Sample | Current and RMS noise at 600 mV | Current / voltage curve (IV curve) |
|---|---|---|---|
| **1** | **3.6 kbp DD with and without nicks** | **9.6 nA, 6.7 pA** | 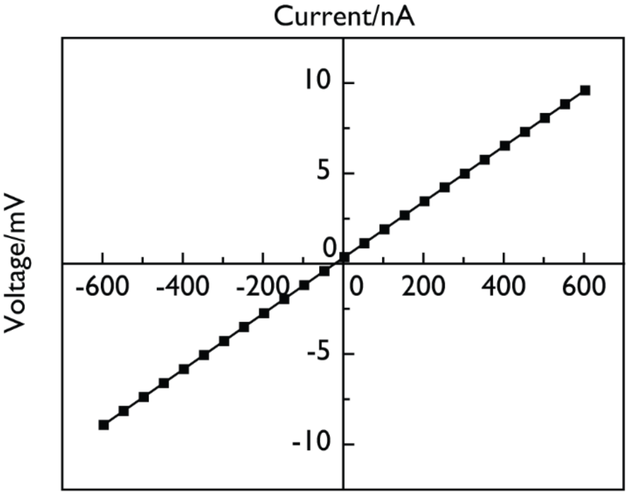 |
| **2** | **3.6 kbp DD with and without nicks** | **12.1 nA, 6.5 pA** | 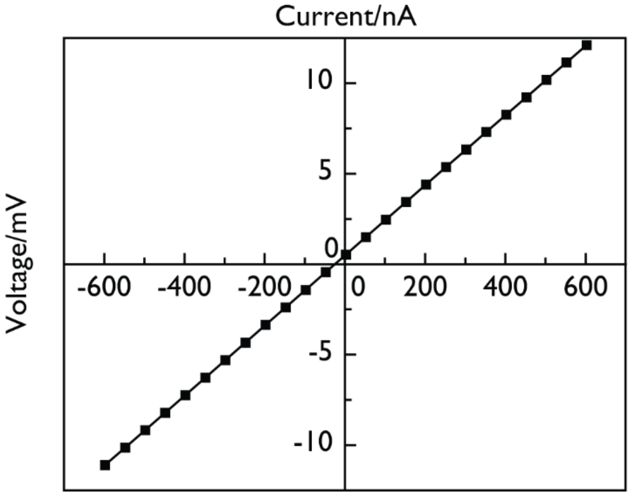 |
